# Extracellular Vesicles from hiPSC-derived NSCs Protect Human Neurons against Aβ-42 Oligomers Induced Neurodegeneration, Mitochondrial Dysfunction and Tau Phosphorylation

**DOI:** 10.1101/2024.07.11.603159

**Authors:** Shama Rao, Leelavathi N Madhu, Roshni Sara Babu, Goutham Shankar, Sanya Kotian, Advaidhaa Nagarajan, Raghavendra Upadhya, Esha Narvekar, James J. Cai, Ashok K Shetty

## Abstract

**Background:** Alzheimer’s disease (AD) is characterized by the accumulation of amyloid beta-42 (Aβ-42) in the brain, causing various adverse effects. Thus, therapies that reduce Aβ-42 toxicity in AD are of great interest. One promising approach is to use extracellular vesicles from human induced pluripotent stem cell-derived neural stem cells (hiPSC-NSC-EVs) because they carry multiple therapeutic miRNAs and proteins capable of protecting neurons against Aβ-42-induced toxicity. Therefore, this *in vitro* study investigated the proficiency of hiPSC-NSC-EVs to protect human neurons from Aβ-42 oligomers (Aβ-42o) induced neurodegeneration.

**Methods:** We isolated hiPSC-NSC-EVs using chromatographic methods and characterized their size, ultrastructure, expression of EV-specific markers and proficiency in getting incorporated into mature human neurons. Next, mature human neurons differentiated from two different hiPSC lines were exposed to 1 µM Aβ-42o alone or with varying concentrations of hiPSC-NSC-EVs. The protective effects of hiPSC-NSC-EVs against Aβ-42o-induced neurodegeneration, oxidative stress, mitochondrial dysfunction, impaired autophagy, and tau phosphorylation were ascertained using multiple measures and one-way ANOVA with Newman-Keuls multiple comparisons post hoc tests.

**Results:** A significant neurodegeneration was observed when human neurons were exposed to Aβ-42o alone. Neurodegeneration was associated with 1) elevated levels of reactive oxygen species (ROS), mitochondrial superoxide, malondialdehyde (MDA) and protein carbonyls (PCs), 2) increased expression of proapoptotic Bax and Bad genes and proteins, and genes encoding mitochondrial complex proteins, 3) diminished mitochondrial membrane potential and mitochondria, 4) reduced expression of the antiapoptotic gene and protein Bcl-2, and autophagy-related proteins, and 5) increased phosphorylation of tau. However, the addition of an optimal dose of hiPSC-NSC-EVs (6 x 10^9^ EVs) to human neuronal cultures exposed to Aβ-42o significantly reduced the extent of neurodegeneration, along with diminished levels of ROS, superoxide, MDA and PCs, normalized expressions of Bax, Bad, and Bcl-2, and autophagy-related proteins, higher mitochondrial membrane potential and mitochondria, enhanced expression of genes linked to mitochondrial complex proteins, and reduced tau phosphorylation.

**Conclusions:** An optimal dose of hiPSC-NSC-EVs could significantly decrease the degeneration of human neurons induced by Aβ-42o. The results support further research into the effectiveness of hiPSC-NSC-EVs in AD, particularly their proficiency in preserving neurons and slowing disease progression.

## Introduction

Alzheimer’s disease (AD) is the fifth leading cause of death, impacting ∼47 million people globally and 7 million people in the United States [1]. It is projected that by 2050, AD-related dementia will affect ∼13 million individuals in the United States [2]. The most prominent pathological features of AD are the accumulation of amyloid plaques in the extracellular space and the neurofibrillary tangles (NFTs) within neurons. These alterations contribute to the progressive loss of synapses, significant chronic neuroinflammation, and eventual neurodegeneration and dementia [3,4]. Amyloid plaques consist of amyloid beta peptide (Aβ), while NFTs contain phosphorylated tau protein [5,6]. The Aβ protein is primarily produced by neurons and, to a lesser extent, by astrocytes in the brain. Additionally, vascular cells and blood cells also generate Aβ [7]. Aβ is created through a series of proteolytic cleavages of amyloid precursor protein (APP) by β- and γ-secretases [8]. After its release, soluble Aβ monomers form dimers, trimers, soluble oligomers, and protofibrils, eventually leading to the formation of fibrils that accumulate in plaques [3]. The main forms of Aβ in the human brain are Aβ1−40 (Aβ-40) and Aβ1−42 (Aβ-42), with Aβ-42 forming aggregates due to being less soluble than Aβ-40 [3,9]. Therefore, Aβ-42 contributes to the pathogenesis of AD and is considered one of the key biomarkers in AD diagnosis [10,11].

While Aβ-42 monomers play roles in neuronal cytoprotective pathways, intracellular signaling, and synaptic functions [12,13], Aβ-42 oligomers (Aβ-42o) can induce neurotoxicity. For example, injection of Aβ-42o into rodent hippocampus can diminish long-term potentiation (LTP), enhance long-term depression (LTD), and cause synapse loss [14–16]. *In vitro* studies using Aβ-42o have reported similar effects on LTP in brain slices [17] and neurotoxicity in rat cortical neurons [18]. Therefore, Aβ-42o pathophysiology seems to be an initial pathological event in AD that triggers subsequent molecular pathways, including tau misfolding, tau buildup in NFTs, and tau-mediated toxicity [11,19–22]. The accumulation of Aβ-42o in the brain leads to various harmful effects. For instance, Aβ-42o can induce increased oxidative stress by inserting into the lipid bilayer, serving as a source of reactive oxygen species (ROS), and initiating lipid peroxidation [23–25]. Additionally, Aβ-42o can localize to mitochondrial membranes, reduce mitochondrial membrane potential, block the transport of nuclear-encoded mitochondrial proteins to mitochondria, disrupt the electron transport chain, increase ROS production, and cause mitochondrial damage [26]. Exposure of neurons to Aβ-42o can also reduce mitochondrial superoxide dismutase 2 and mitochondrial cytochrome C protein [27]. Furthermore, mitochondrial-specific accumulation of Aβ-42o induces mitochondrial dysfunction leading to apoptotic cell death [28]. Apoptotic cell death in response to Aβ-42o exposure involves altered expression of pro- and anti-apoptotic proteins [29]. Furthermore, Aβ-42o exposure to neurons can significantly increase the phosphorylation of tau protein by activating tau kinases, mitogen-activated protein kinase, and glycogen synthase kinase-3 beta [30,31]. Therefore, there is a keen interest in therapeutic interventions aimed at reducing Aβ-42o-induced toxicity in AD.

One potential approach for safeguarding neurons from Aβ-42o induced toxicity in AD is using extracellular vesicles (EVs) released from stem cells, as these EVs contain various therapeutic miRNAs and proteins that can protect neurons [32–35]. However, no previous studies have tested the effectiveness of neural stem cell (NSC)-derived EVs in amyloid toxicity models. Recent studies in our laboratory have demonstrated that EVs released by human induced pluripotent stem cell (hiPSC)- derived NSCs (hiPSC-NSC-EVs) are enriched with miRNAs and proteins that can mediate antioxidant, antiapoptotic, anti-inflammatory, and neurogenic effects [36–38]. Furthermore, hiPSC-NSC-EVs contain specific miRNAs (miRNA-103a-3p, miRNA-21-5p) and proteins (pentraxin-3) that can promote neuroprotection, including through antiapoptotic mechanisms [39–42]. These EVs are also enriched with miR-148a-3p, which can inhibit the phosphorylation of tau by directly targeting p35/cyclin dependent kinase 5 and phosphatase and tensin homolog (PTEN)/p38 mitogen-activated protein kinase (MAPK) pathways [43]. Additional protein (e.g., hemopexin) enriched in hiPSC-NSC-EVs can maintain mitochondrial function and homeostasis and reduce ROS toxicity [36,44]. Moreover, the absence of tumorigenic and immunogenic properties, combined with the capacity to efficiently penetrate the entire brain and integrate into neurons following intranasal administrations [36,45] positions hiPSC-NSC-EVs as a compelling biologic for conferring neuroprotection in neurodegenerative disorders. Therefore, the current *in vitro* study examined whether hiPSC-NSC-EVs could protect human neurons derived from two different hiPSC lines against Aβ-42o-induced neurodegeneration. The results provide novel evidence that an optimal dosage of hiPSC-NSC-EVs could markedly avert Aβ-42o-induced neurodegeneration by 1) maintaining higher mitochondrial membrane potential, 2) reducing total ROS and mitochondria-generated superoxide production, 3) attenuating mitochondrial hyperactivation and loss, 4) modulating pro-apoptotic and antiapoptotic gene and protein expression, 5) regulating autophagy, and 6) mitigating tau phosphorylation.

## Materials and Methods

### Differentiation of hiPSC-NSCs into neurons

The mature human neurons used in these studies were freshly differentiated from passage 10 human NSCs (hNSCs) generated from two hiPSC lines: one from Wisconsin International Stem Cell Bank (IMR90-4) and the other from the National Institutes of Health (NIH; LiPSC-GR1.1, Research grade). The hiPSCs were first transformed into NSCs using a directed differentiation protocol, as described in our previous studies [36,37]. Then, passage 10 hNSCs were used to obtain mature human neurons, using the protocol published at the Thermo Fisher Scientific website with minor modifications. Briefly, a homogenous population of passage 10 NSCs were seeded onto 24-well plates coated with poly-D-lysine (0.1 mg/ml, Sigma Aldrich, St. Louis, MO, USA), poly-L-ornithine (0.1 mg/ml, Sigma Aldrich) and 10 µg/ml laminin (Corning Inc., Corning, NY, USA) at a density of 15,000 cells/cm^2^ using neuronal differentiation media. The media comprised Neurobasal (1X, Thermo Fisher Scientific), GlutaMAX-I (2 mM, Thermo Fisher Scientific), serum-free B27 supplement with retinoic acid (2%, Thermo Fisher Scientific) and brain-derived neurotrophic factor (BDNF, 10 ng/mL, Thermo Fisher Scientific). The cultures were maintained for 30 days with media exchange every 3 days to enable complete differentiation of hNSCs into neurons. The neuronal differentiation was confirmed through immunofluorescence staining, using antibodies against microtubule-associated protein-1 (rabbit anti-Map-2; Millipore, Burlington, Massachusetts, USA) and neuron-specific nuclear antigen (rabbit anti-NeuN; Millipore). The percentage of neurons among all cells was quantified from cultures processed for Map-2 immunofluorescence with DAPI counterstaining (6 independent cultures, 3 images per culture, total 18 images).

### Isolation hiPSC-NSC-EVs via chromatographic methods

After reaching 80% confluency, passage 10 hiPSC-NSCs were dislodged using TrypLE^TM^ Express enzyme (Thermo Fisher Scientific, Waltham, MA, USA), washed with NSC expansion medium containing equal volumes of Neurobasal media and DMEM/F-12 (Thermo Fisher Scientific), Neural Induction supplement (1% of media, Thermo Fisher Scientific) and penicillin-streptomycin (100 units, ThermoFischer Scientific), and then seeded at 1000 cells/cm^2^ onto T300 tissue culture flasks (VWR, Radnor, Pennsylvania, USA) containing fresh NSC expansion medium. The NSCs were maintained for 7 days, and spent media were harvested twice on day 4 and 7. The EVs were isolated using anion-exchange chromatography (AEC) followed by size exclusion chromatography (SEC). The SEC fractions containing a higher number of EVs with lower protein content were combined, concentrated, and stored at −20°C for future use.

### Characterization of hiPSC-NSC-EVs for EV-specific marker expression

We used nanoparticle tracking analysis to determine the concentration and size of EVs, as outlined in our previous studies [36,37]. We assessed the presence of EV-specific tetraspanin markers CD63 and CD81, intra-EV marker protein ALIX, and a non-EV marker cytochrome C using western blotting. In brief, we mixed 25 x 10^9^ EVs with 100 µl of a mammalian protein extractor reagent (Thermo Fischer Scientific), lysed it as previously described [46], and the total protein concentration was determined using the Pierce BCA protein assay kit (ThermoFisher Scientific). Next, we loaded and separated 40 µg of protein using 4-12% NuPAGE Bis-Tris Gels (ThermoFisher Scientific). The proteins were then transferred onto nitrocellulose membranes using ThermoFisher Scientific’s iBlot2 gel transfer technology. We processed the membrane for protein detection using antibodies against CD63 (1:1000; BD Biosciences, San Jose, California, USA), CD81 (1:1000; BD Biosciences), and ALIX (1:1000; Santa Cruz Biotechnology, Dallas, TX, USA). The chemiluminescent signal was detected using ibright 1500 (Thermo Fischer Scientific) and an ECL kit (Thermo Fischer Scientific). In addition, we examined the expression of cytochrome C to check for contamination of EV preparations with deep cellular proteins compared to the hNSC lysate [36].

### Visualization of the morphology of hiPSC-NSC-EVs through transmission electron microscopy

We used transmission electron microscopy (TEM) to examine the ultrastructure of hiPSC-NSC-EVs, following the methods outlined in our previous studies [36,47]. To do this, we placed 5 μl (∼1.5 x 10^9^ EVs) of the EV suspension, diluted in PBS, on 300 mesh carbon-coated copper grids (Electron Microscopy Sciences, Hatfield, PA, USA) at room temperature. After five minutes, we blotted the excess liquid from the grids with filter paper and rinsed them twice with distilled water. We then stained the grids by continuously dripping a 150 μl solution of 0.5% uranyl acetate at a 45° inclination. After removing the excess liquid, we air-dried the grids at room temperature for 10 minutes. The images were captured using an FEI Morgagni 268 transmission electron microscope with a MegaView III CCD camera. Lastly, we computed the diameters of hNSC-EVs using the “Analyze” tool in the ImageJ software, averaging the measured diameters along four axes (x, y, x + 45°, y + 45°).

### Evaluation of the neuroprotective effects of hiPSC-NSC-EVs on neurons exposed to Aβ-42o

First, 30-day-old human neuronal cultures were exposed to various doses of Aβ-42o (Sigma-Aldrich, St. Louis, Missouri, USA), ranging from 0.1 µM to 5 µM, for 24 hours to determine the dose-dependent cytotoxicity using the MTT assay. For subsequent studies, 30-day-old human neuronal cultures were exposed to 1 μM Aβ-42o and were treated concurrently with hiPSC-NSC-EVs at three different concentrations (1.5, 3.0, or 6 x10^9^, (3 independent cultures/group). Additionally, separate sets of control sister cultures with or without Aβ-42o exposure were maintained (3 biological replicates/group). Following a 24-hour incubation period, the neuronal cultures were utilized for various assays.

### Visualization of the interaction of cultured human neurons exposed to Aβ-42o with hiPSC-NSC-EVs

The hiPSC-NSC-EVs were labeled with the PKH26 dye using the PKH26 Red Fluorescent Cell Linker Kit for general cell membrane labeling (Sigma-Aldrich). Approximately 30 x 10^9^ EVs were mixed with 100 µl of PBS without calcium and magnesium, followed by the addition of 900 µl of diluent C. Then, 4 µl of PKH26 dye was added to the mixture to achieve a final concentration of 8 μM (dye solution). After 10 minutes of incubation in the dark, the 1 ml preparation was diluted to a final volume of 5 ml using PBS without calcium and magnesium, then centrifuged using ultrafiltration columns (100 kDa, Corning Inc.). This washing step was repeated four times to remove the excess dye from the EV preparation. These EVs (1X10^9^) were introduced to mature human neuronal cultures with or without 1 μM Aβ-42o (3 independent cultures/group), and the cultures were fixed in 2% paraformaldehyde after 4 hours of incubation. The cultures were then processed for MAP-2 immunofluorescence to observe EVs contacting neurons, including their internalization by neurons. The interactions were photographed using a fluorescent microscope.

### Validation of selected miRNAs and pentraxin 3 in hiPSC-NSC-EVs

Our previous small RNA sequencing and proteomics studies have demonstrated enrichment of hiPSC-NSC-EVs with miR-148a-3p, miR-21-5p, miR-103a-3p and pentraxin 3 (PTX3), in addition to multiple other miRNAs and proteins [36]. Because of their proficiency in mediating antiapoptotic effects (miR-21-5p, miR-103a-3p, and PTX3) [42,48–55], and inhibiting tau phosphorylation (miR-148a-3p) [43], we confirmed their expression in hiPSC-NSC-EVs employed in the current study. For validating the enrichment of miR-148a-3p, miR-21-5p, and miR-103a-3p in hiPSC-NSC-EVs, the total RNA was isolated from two biological replicates of hiPSC-NSC-EVs (∼10 × 10^9^ each) using the SeraMir Exosome RNA amplification kit (System biosciences, Palo Alto, CA, USA). miRCURY LNA RT Kit (Qiagen, Germantown, MD, USA) was employed for converting 5 ng/μl of total RNA into cDNA. miRCURY LNA miRNA SYBR Green PCR kit (Qiagen) and miRCURY LNA miRNA PCR assay primer mix (Qiagen) were used to measure the comparative expression of these miRNAs (two technical replicates/biological replicate). We employed ELISA to confirm the enriched expression of PTX3 in hiPSC-NSC-EVs. Two biological replicates of hiPSC-NSC-EVs (∼50 × 10^9^ each) were lysed, and PTX3 was measured using a commercially available ELISA kit (Aviscera Biosciences, Sunnyvale, CA, USA). The concentration of PTX3 was normalized to the total protein content in the EV lysate.

### Cytotoxicity measurement by MTT assay

After incubating human neuronal cultures with Aβ-42o or Aβ-42o and hiPSC-NSC-EVs for 24 hours, we added 300 µl of MTT solution (0.5 mg/ml, Sigma-Aldrich) to each culture well and then incubated them for 4 hours at 37°C. The wells were then treated with 300 µl of DMSO to release the purple formazan into the solution, and we recorded the absorbance at 570 nm (3-5 technical replicates for each of 3 biological replicates/group). We calculated the percentage of cell viability and compared it across the study groups.

### Measurement of human neuronal viability via live and dead cell assay

Neuronal cultures were first incubated with Aβ-42o alone or Aβ-42o with different doses of hiPSC-NSC-EVs for 24 hours (3 biological replicates/group). Then, we employed the ReadyProbesTM Cell Viability Imaging Kit (Invitrogen, Waltham, MA, USA), which identified live cells as blue and dead cells as green in neuronal cultures. We added NucBlueTM Live Reagent and NucGreenTM Dead Reagent (two drops each for 1 ml of neuronal media) and incubated cultures in these reagents for 30 minutes at 37°C. Next, the fluorescent images were randomly taken from live cultures using a Nikon inverted fluorescence microscope. We quantified the extent of dead cells by measuring the area fraction of green fluorescence in images using Image J (3 images/culture, a total of 9 images/group).

### Measurement of the survival of Map-2 positive neurons

The cultures from all groups (control, Aβ-42o, and Aβ-42o+hiPSC-NSC-EVs groups) were processed for Map-2 immunofluorescence staining using an antibody against Map-2 (ThermoFisher Scientific) and counterstained with DAPI (ThermoFisher Scientific) to assess the number of surviving neurons present in different cultures (3 independent cultures/group). Then, neuronal counts per 0.35 mm^2^ unit area were taken from randomly chosen 9 images from 3 independent cultures/group.

### Quantification of oxidative stress markers

Three independent neuronal cultures from each group were separately lysed in ice-cold RIPA buffer (Life Technologies, Carlsbad, CA, USA) with protease and phosphatase inhibitor cocktail (Life Technologies) using sonication for 10 seconds. The lysed solution was centrifuged twice, and aliquots of the supernatant solution were stored at -80°C until further use. The protein concentration in different samples was measured using a Pierce BCA reagent kit (ThermoFisher Scientific). We followed the manufacturer’s instructions for measuring malondialdehyde (MDA Cayman Chemical; Ann Arbor, MI, USA) and protein carbonyls (PCs, Cayman Chemical) from neuronal culture lysates (2 technical replicates from each of 3 biological replicates).

### Measurement of the expression of genes related to oxidative stress and apoptosis in neuronal cultures

We conducted qRT-PCR to measure the expression of genes involved in regulating oxidative stress (iNos and Cox2) and apoptosis (Bad, Bax, and Bcl2) in neuronal cultures from all groups (control, Aβ-42o and Aβ-42o+hiPSC-NSC-EVs groups, 3 independent cultures/group). Total RNA was extracted using the RNeasy kit (Qiagen, Valencia, CA, USA), as outlined in our previous reports [56]. The extracted RNA (500 ng) was then converted to cDNA using the RT2 First Strand Kit (Qiagen) and stored at −20°C until further analysis [56]. The reactions were carried out according to the manufacturer’s protocol using a CFX96 Real-Time System (Bio-Rad, Hercules, CA, USA). PCR amplification and melt curve analysis were performed as detailed in our previous report [56]. Finally, the 2^dd-Ct values for each gene were compared across different groups.

### Measurement of total ROS from human neuronal cultures

The hNSC-derived neurons were cultured in 24 well-plates, and 30-day-old neuronal cultures were randomly assigned to each of the three groups (control, Aβ-42o or Aβ-42o+EVs groups, 3 biological replicates/group). Then, the cultures assigned to Aβ-42o or Aβ-42o+EVs groups were incubated with Aβ-42o alone or Aβ-42o with hiPSC-NSC-EVs for 24 hours. Next, following three washes in Dulbecco’s phosphate buffered saline (DPBS), cultures from all groups were incubated with 10 mM DCFH-DA solution (Sigma Aldrich) prepared in a prewarmed neuronal differentiation medium for 30 minutes [57]. The cultures were again washed 3 times with DPBS and observed under a Nikon fluorescence microscope. After confirming the appearance of green fluorescence, the cells in each culture were lysed with radioimmunoprecipitation assay (RIPA) buffer. After 5 minutes, the lysates were collected and centrifuged at 14,000 rpm for 10 minutes. The supernatant was next transferred to black 96-well pates, and fluorescence intensity was measured in a fluorometer (TECAN). Two technical replicates per each of three biological replicates/group were employed.

### Measurement of mitochondrial superoxide (MitoSOX)

We employed an assay using MitoSOX green (Thermo Fisher Scientific) to detect the extent of mitochondria-generated superoxide in human neuronal cultures. MitoSOX green can enter mitochondria, where it can be oxidized only by superoxide and not by other ROS or reactive nitrogen species. The oxidized green binds to nucleic acids in mitochondria and produces a green fluorescence [58]. Hence, the application of MitoSOX green facilitates the detection of the extent of mitochondria-generated superoxide in live neuronal cultures. We prepared 1 µM MitoSox green using Hanks balanced salt solution (HBSS, Gibco) and incubated cultures from all three groups (control, Aβ-42o or Aβ-42o+EVs groups, 3 biological replicates/group) in this solution for 30 minutes at standard culture conditions. Afterward, the cultures were observed under a Nikon fluorescence microscope to detect green fluorescence emission, and plates were immediately read in a fluorometer (TECAN).

### Quantification of proteins levels of BCL-2 and BAD by ELISA

The selected antiapoptotic and proapoptotic markers BCL2 and BAD levels were measured in neuronal cultures from all groups (control, Aβ-42o, and Aβ-42o+hiPSC-NSC-EVs groups, 3 biological replicates/group). We followed the manufacturer’s protocol to measure the protein concentrations of BCL2 (Invitrogen) and BAD (MyBioSource, San Diego, CA, USA) in the cell lysates of neuronal cell cultures.

### Measurement of mitochondrial respiration complex genes

We evaluated the expression of genes linked to different mitochondrial complexes, including complex I (Ndufs6, Ndufs7), complex II (Sdha, Sdhb), complex III (Cyc1, Bcs1l), complex IV (Cox6a1), and complex V (Atp6ap1) in RNA samples extracted from neuronal cultures belonging to different experimental groups (control, Aβ-42o, and Aβ-42o+hiPSC-NSC-EVs, 3 biological replicates/group). We used specific primers (Genecopoeia, Rockville, MD, USA) to amplify the cDNA samples and then normalized the obtained CT values with the housekeeping gene Gapdh. The 2^dd-Ct values for each gene were then compared across the different groups of neuronal cultures.

### Measurement of mitochondrial membrane potential using JC-1 dye

An assay utilizing a dye called JC-1 was used to determine changes in the mitochondrial membrane potential in live human neurons [59]. This assay provides a high ratio of red fluorescence when the mitochondrial potential is high (i.e., healthy mitochondria) and a high ratio of green fluorescence when the mitochondrial potential is low (i.e., damaged mitochondria), facilitating the quantification of the health of neuronal mitochondria. The mature neuronal cultures from all three groups (control, Aβ-42o or Aβ-42o+ 6x10^9^ EVs groups, 3 biological replicates/group) were incubated with JC-1 solution (Sigma Aldrich) for 20 minutes at 37°C in the dark. The green fluorescence intensity and red fluorescence intensity were measured using a Spark control microplate spectrofluorometer (TECAN). Another set of parallel cultures were washed in DPBS after incubation in JC-1 solution and images were taken from randomly chosen areas in each culture using a 20X objective lens in a Nikon Inverted fluorescence microscope. Afterward, the AF of red fluorescence in each image was measured using Image J (3 images per culture, total 9 images/group).

### Measurement of total mitochondria per individual neuron

The mature neuronal cultures from all three groups (control, Aβ-42o or Aβ-42o+EVs groups, 3 biological replicates/group) were fixed in 2% paraformaldehyde and processed for dual immunofluorescence staining using antibodies against TOM20 (a mitochondria marker, ProteinTech, Rosemont, IL, USA) and Map-2 (a neuronal marker, ThermoFisher Scientific). After counterstaining with DAPI (ThermoFisher Scientific), we visualized TOM20+ mitochondria within Map-2+ neurons in a Nikon inverted microscope, and images were taken from randomly chosen areas in each culture using a 20x objective lens. By using ImageJ, the total number of neurons and areas occupied by neurons (i.e., Map-2+ structures) and mitochondria (i.e., TOM20+ structures within neurons) were measured in each image (3 images per culture, a total of 9 images/group). Next, the AF of TOM20+ structures for all neurons in each image was subtracted by the number of neurons to calculate the average AF of TOM20+ structures per individual neuron in each image.

### Measurement of autophagy related protein, beclin1 and MAP1LC3B

Autophagy is a fundamental process that clears the accumulation and deposition of toxic proteins in the cellular system [60]. We measured autophagy-related markers such as beclin1 (MyBiosource) and MAP1LC3B (MyBiosource) in neuronal cultures belonging to different experimental groups (control, Aβ-42o, and Aβ-42o+hiPSC-NSC-EVs, 3 biological replicates/group).

### Quantification of total tau and phospho-tau

We first performed a standard sandwich ELISA as per the manufacturer’s guidelines for the measurement of total tau (Invitrogen) and phospho-tau (Invitrogen) in neuronal cultures belonging to different experimental groups (control, Aβ-42o or Aβo+EVs groups, 3 biological replicates/group). The concentrations of these proteins were normalized to 1 mg of total protein in the cell lysate. Next, we performed western blots for total tau and phospho-tau to confirm ELISA results (3 biological replicates/group). We loaded 30 μg of protein onto 4–12% NuPAGE Bis-Tris Gels (ThermoFisher Scientific) and separated. After transferring proteins onto a nitrocellulose membrane using the iBlot2 gel transfer device (ThermoFisher Scientific), proteins in the membrane were detected using antibodies against tau (1:200, Santa Cruz Biotechnology, Dallas, Texas, USA), p-tau (1:1000, ThermoFisher Scientific) and GAPDH (1:5000, Millipore). GAPDH was used as a loading control for cell lysates. Then, the protein signals were detected using the chemiluminescence reagents provided with the ECL detection kit (ThermoFisher Scientific) and visualized using the iBright imaging system (ThermoFisher Scientific). ImageJ software was used to determine the intensity of each targeted band by normalizing it to a corresponding GAPDH band.

### Statistical Analyses

The experiments utilized 3 independent cultures per condition/group, and statistical analysis was done using Prism software 10.0. A one-way ANOVA with Newman-Keuls multiple comparison post-hoc tests was employed for datasets containing three or more groups comparison. In all comparisons, p<0.05 was considered a statistically significant value.

## Results

### Establishment of hiPSC-NSCs and primary human neuronal cultures from hiPSC-NSCs

Incubation of the hiPSC colony (Fig. 1 [A]) with the neurobasal medium containing neural induction supplement for nine days resulted in the generation of cells having small polygonal to spindle-shaped morphology. Subsequent serial passaging of these cells eliminated undifferentiated hiPSCs, and all cells in passage 11 cultures expressed NSC markers Sox-2 and nestin (Fig. 1 [B-D]). Furthermore, passage 10 hNSCs from two different hiPSC lines subjected to neuronal differentiation protocol gave rise to predominantly mature neurons expressing markers such as Map-2 and NeuN (Fig. 1 [E-L]). The percentage of mature neurons in these cultures was 84.72±4.52%. From here onwards, the neurons derived from two distinct hiPSC lines are referred to as iNeurons 1 and iNeurons 2.

**Figure 1:**
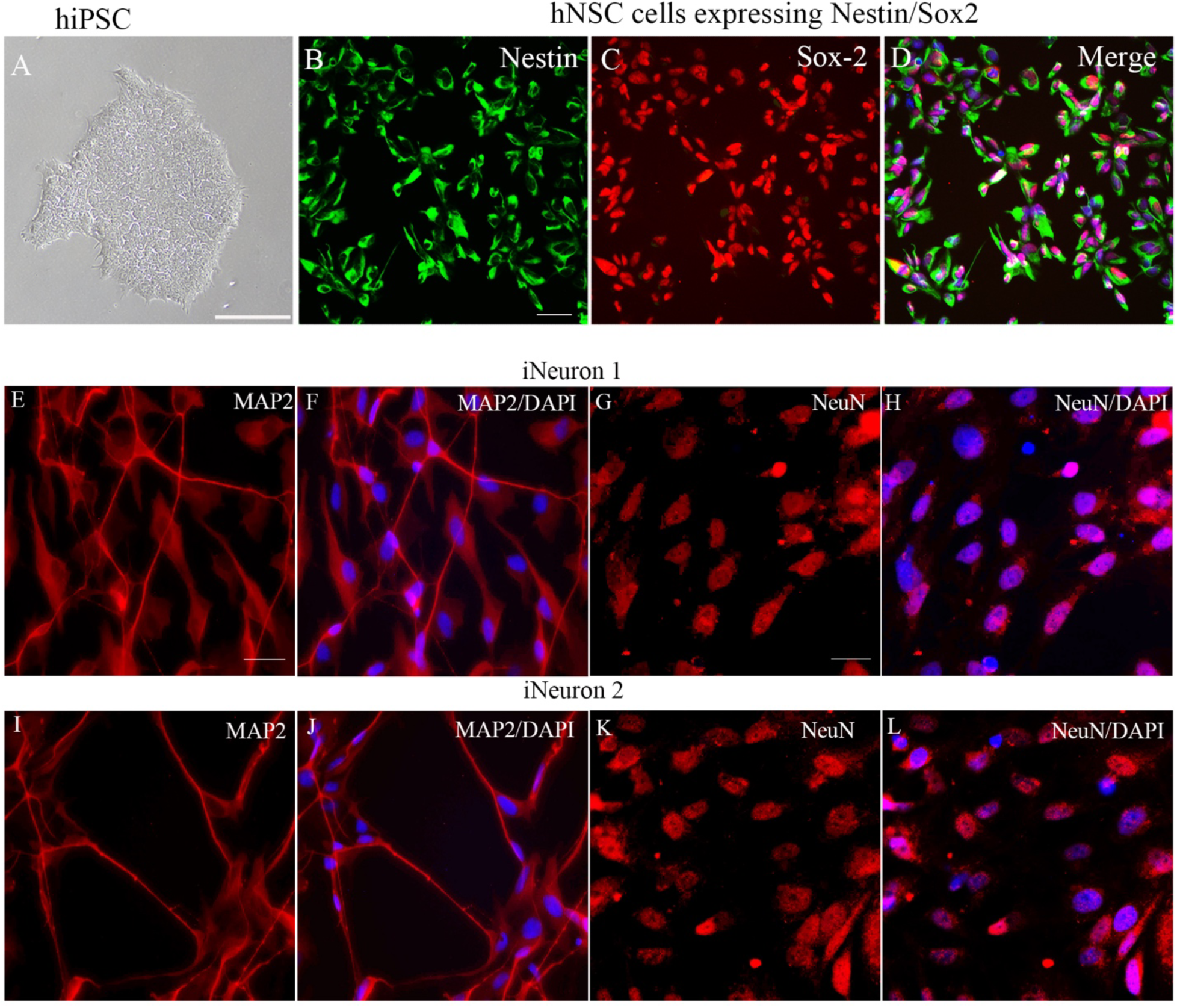
Generation and characterization of human neural stem cells (NSCs) and human neurons from human induced pluripotent stem cells (hiPSCs). Figure A shows a hiPSC colony. Images B-D show the expression of nestin (B), Sox-2 (C), and both nestin and Sox-2 counterstained with DAPI (D) in NSCs derived from hiPSCs. Figures E-H and I-L respectively illustrate mature neurons derived hiPSC lines 1 and 2 (iNeurons 1 and 2) immunostained with microtubule-associated protein-2 (MAP-2; E-F, I-J) and neuron-specific nuclear antigen (NeuN; G-H, K-L). Scale bars: 100 µm.

### hiPSC-NSC-EVs exhibited a characteristic ultrastructure and expressed CD63, CD81, and Alix

The SEC fractions 7–13 contained considerably higher numbers of EVs with minimal co-eluting proteins (Fig. 2 [A]); hence, only the fractions 7-13 were pooled and used for further studies. Nanoparticle tracking analysis showed that the size of EVs ranged from 100 to 200 nm with a mean value of 121.6 ± 0.6 nm (Fig. 2 [B]). Imaging with a TEM demonstrated round or oval vesicles with lipid bilayer membranes ranging from 50 to 130 nm in diameter. Examples of individual EVs exhibiting different sizes and shapes found in our hiPSC-NSC-EVs preparations are illustrated in Figure 2 (Fig. 2 [C]). Additionally, western blot analysis showed that EVs expressed several EV markers, including CD63, CD81, and ALIX (Fig. 2 [D]). The EV preparation lacked the deep cellular protein such as the cytochrome C (Fig. 2 [D]).

**Figure 2:**
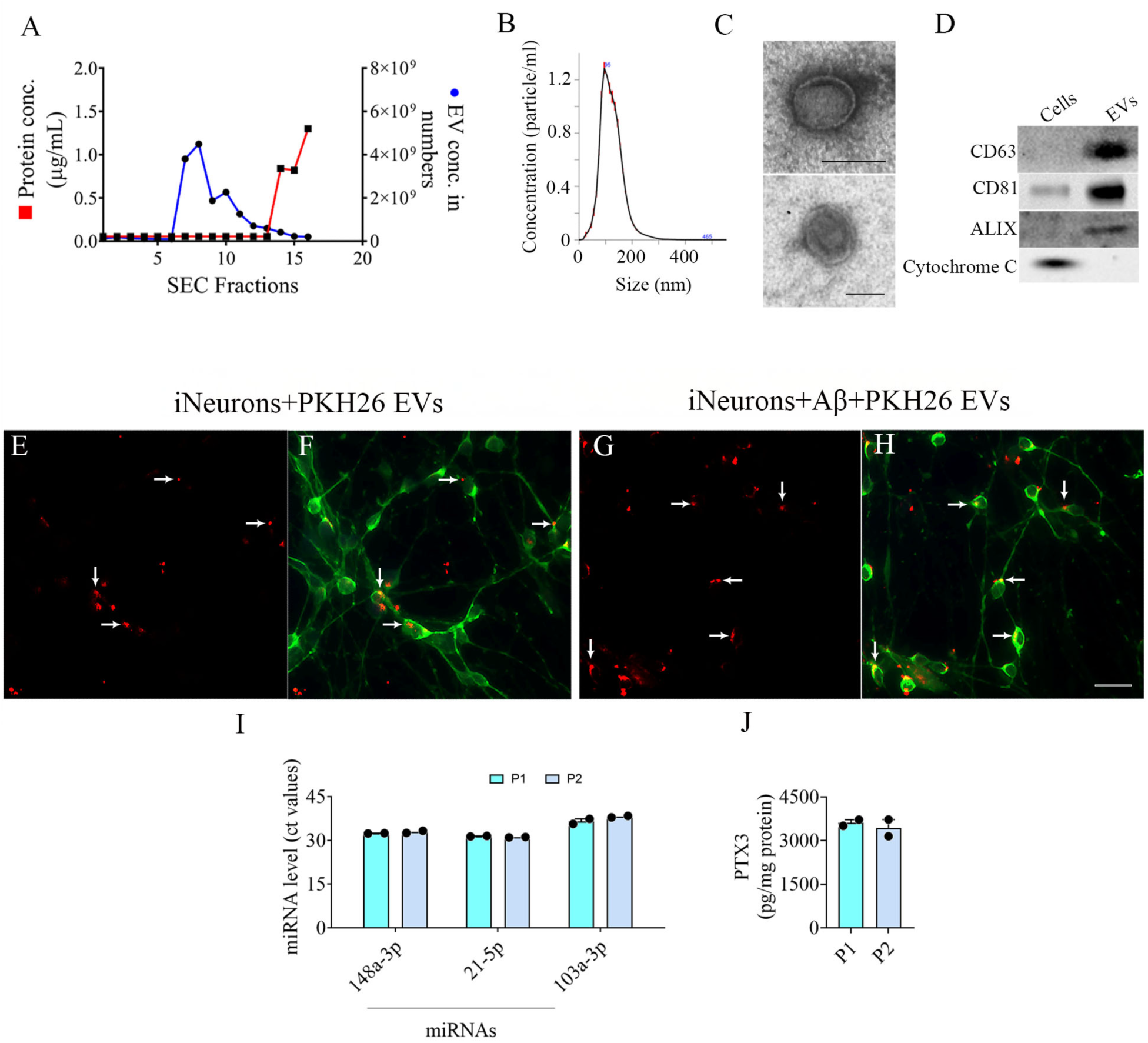
Characterization of extracellular vesicles from human induced pluripotent stem cell-derived neural stem cells (hiPSC-NSC-EVs). Graph A illustrates the number of EVs vis-a-vis protein content in various fractions collected from size-exclusion chromatography. Graph B shows that the average size of EVs varied between 100-200 nm, with a mean size of 121.6 nm. Figure C illustrates the morphology of EVs examined through transmission electron microscopy. Scale bar, 100 nm. Figure D shows the expression of CD63, CD81, and ALIX but not cytochrome C in hiPSC-NSC-EVs visualized through Western blotting. Full-length blots of CD63, CD81, ALIX and cytochrome C are presented in Supplementary figures 1-4. Figures E-H demonstrate the incorporation of PKH26-labeled EVs (red particles) into microtubule-associated protein-2 (MAP-2)-positive human neurons (green) in standard cultures (E-F) and Aβ-42 oligomers exposed cultures (G-H). Scale bar, 100 µm. The bar charts show Ct values of different miRNAs measured via qPCR (I) and pentraxin 3 (PTX3) measured via ELISA (J) from two biological replicates of hiPSC-NSC-EVs.

### Visualization of the interaction of PKH26-labelled hiPSC-NSC-EVs with mature human neurons exposed to *Aβ-42o*

The experiment involved the addition of PKH26-labeled hiPSC-NSC-EVs to two types of neuronal cultures: one was standard cultures, and the other was exposed to Aβ-42o. The results demonstrated that hiPSC-NSC-EVs could interact with neurons in both cultures. Whether in a standard or adverse environment, the hiPSC-NSC-EVs were observed to either contact the outer membranes of neurons or to be taken up by the neurons themselves, as depicted in Fig. 2 [E-H]. Notably, the presence of Aβ-42o did not disrupt the ability of hiPSC-NSC-EVs to interact with the neurons.

### Validation of miR-148a-3p, miR-21-5p, miR-103a-3p and PTX3 in hiPSC-NSC-EVs

qPCR analysis confirmed the presence of miRNAs such as miR-148a-3p, miR-21-5p, miR-103a-3p in two biological replicates hiPSC-NSC-EVs employed in the study (Fig. 2 [I]). Similarly, ELISA confirmed the enrichment of PTX3 in two biological replicates of hiPSC-NSC-EVs used in the study (Fig. 2 [J]).

### Aβ-42o induced neurodegeneration in hiPSC-NSC-derived neurons is dose-dependent

The exposure of mature neurons to different concentrations of Aβ-42o induced significant neurodegeneration (Fig. 3 [A]). Measurement of the percentage neuronal viability using an MTT assay indicated that Aβ-42o induced neurodegeneration is dose-dependent with a lower dose (0.1 µM) causing a moderate decrease in viability (p<0.05, Fig. 3 [A]), whereas higher doses (0.5, 1.0, 1.5 or 5 µM) causing much severe, progressive decreases in neuronal viability (p<0.0001, Fig. 3 [A]). Exposure to Aβ-42o at 5 µM concentration reduced viability to ∼53% of the control group (Fig. 3 [A]). Thus, the subsequent experiments were performed using the sublethal dose of 1 µM Aβ-42o.

**Figure 3:**
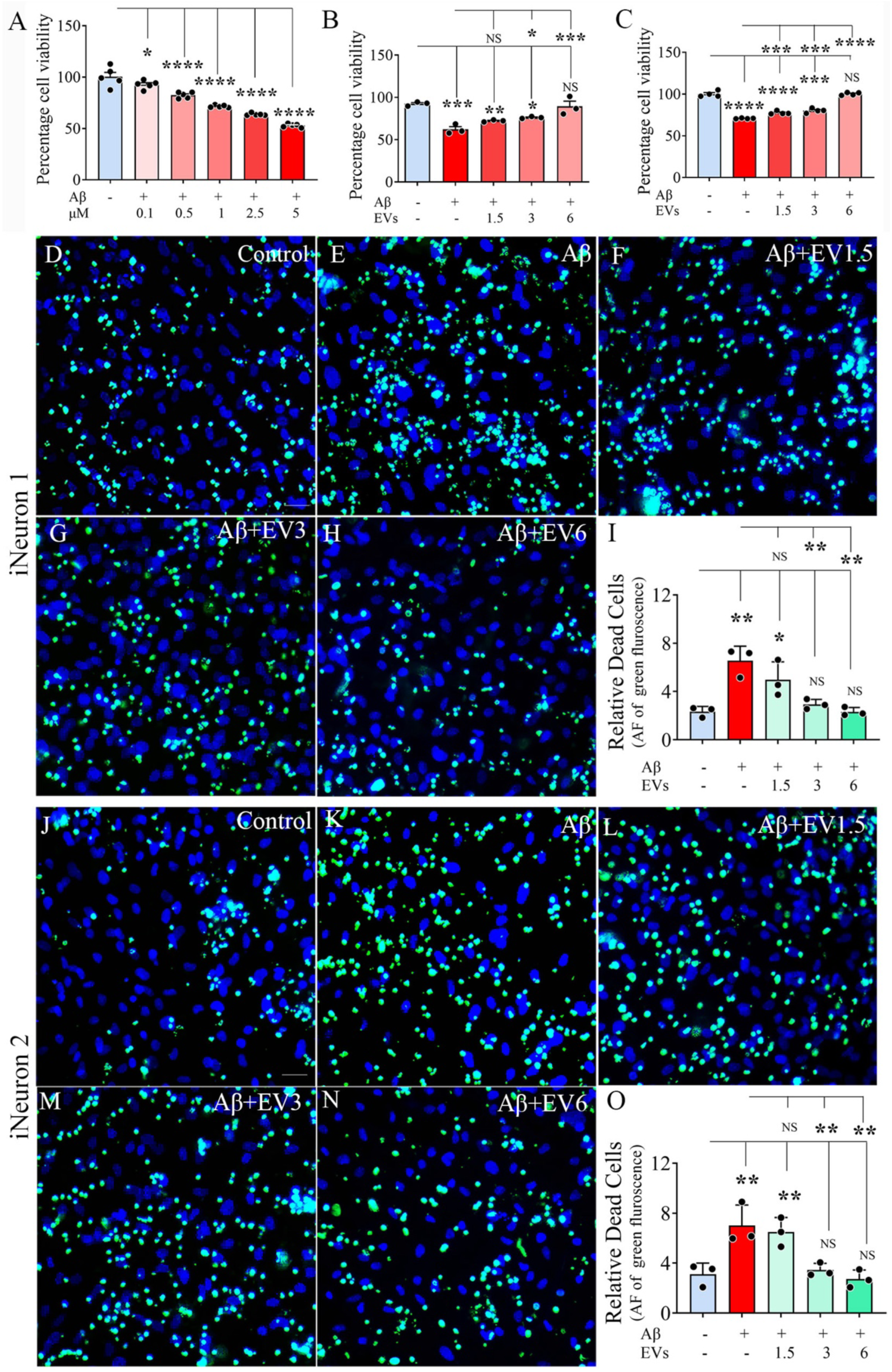
Assessment of cell viability in iNeurons cultures exposed to Aβ-42 oligomers (Aβ-42o), or Aβ-42o and extracellular vesicles from human induced pluripotent stem cell-derived neural stem cells (hiPSC-NSC-EVs). The bar chart in A shows the results of an MTT assay comparing the percentages of viable neurons in iNeurons cultures exposed to different concentrations of Aβ-42o. The bar charts B-C show the results of MTT assays comparing the percentages of viable neurons in iNeurons 1 and 2 cultures exposed to Aβ-42o or Aβ-42o and different concentrations of hiPSC-NSC-EVs. Figures D-H and J-N are representative images from cultures of iNeurons 1 (D-H) and iNeurons 2 (J-N), processed for live and dead cell assay. D and J, control iNeurons 1 and 2 cultures. E and K, iNeurons 1 and 2 cultures exposed to Aβ-42o. F and L, iNeurons 1 and 2 cultures exposed to Aβ-42o and treated with 1.5 x 10^9^ billion EVs. G and M, iNeurons 1 and 2 cultures exposed to Aβ-42o and treated with 3.0 x 10^9^ billion EVs. H and N, iNeurons 1 and 2 cultures exposed to Aβ-42o and treated with 6.0 x 10^9^ billion EVs. The bar charts I and O compare the area fractions of green fluorescence (implying dead cells) across different groups for iNeurons 1 (I) and iNeurons 2 (O) cultures; Scale bar, 100 µm.

### hiPSC-NSC-EVs protected human neurons against Aβ-42o induced neurodegeneration

Increasing doses (1.5, 3, or 6 x 10^9^) of hiPSC-NSC-EVs to Aβ-42o exposed human neuronal cultures reduced neurodegeneration. The protective effects were dose-dependent in both iNeurons 1 and iNeurons 2 cultures. In iNeuron 1 cultures, compared to cultures receiving Aβ-42o alone, Aβ-42o exposed neuronal cultures receiving hiPSC-NSC-EVs treatment at 3 or 6 x 10^9^ dose displayed better survival of neurons (p<0.05-0.001, Fig. 3 [B]). In iNeurons 2 cultures, hiPSC-NSC-EVs treatment at 1.5, 3, or 6 x 10^9^ doses considerably improved the survival of neurons (p<0.001-0.0001, Fig. 3 [C]). Furthermore, fluorescence imaging from the live and dead cell assay (Fig. 3 [D-H, J-N]) revealed significantly higher green fluorescence (implying increased cell death) in the Aβ-42o exposure groups compared to control neuronal cultures in both iNeurons 1 and iNeurons 2 cultures (Fig. 3 [D-E, J-K]). AF analysis of green fluorescence using image J from iNeurons 1 and iNeurons 2 cultures demonstrated significantly higher values in Aβ-42o alone group compared to control (p<0.01, Fig. 3 [I, O]). Interestingly, in Aβ-42o exposed neuronal cultures receiving 3 or 6 x 10^9^ hiPSC-NSC-EVs, AFs of green fluorescence were reduced compared to Aβ-42o alone group (p<0.01, Fig. 3 [I, O]). Moreover, these values matched values in the corresponding control neuronal cultures (p<0.01, Fig. 3 [I, O]).

To further confirm the above findings, we quantified the surviving Map2+ neurons (i.e., number of neurons per unit area) in images taken randomly from different neuronal culture groups (Fig. 4 [A-E, G-K]). In both iNeurons 1 and iNeurons 2 cultures, the survival was diminished in all Aβ-42o exposed groups compared to the control group (p<0.0001, Fig. 4 [F, L]). However, compared to neuronal cultures exposed to Aβ-42o alone, Aβ-42o exposed neuronal cultures receiving different doses of hiPSC-NSC-EVs displayed increased survival of neurons in a dose-dependent manner (Fig. 4 [F, L]). In iNeurons 1 cultures, compared to neuronal cultures exposed to Aβ-42o alone, cultures receiving hiPSC-NSC-EVs treatment at 3 or 6 x 10^9^ doses displayed better survival of neurons (p<0.05-0.0001, Fig. 4 [F]). In iNeurons 2 cultures, hiPSC-NSC-EVs treatment at 1.5, 3, or 6 x 10^9^ doses considerably improved the survival of neurons (p<0.05-0.0001, Fig. 4 [L]). Thus, multiple measures of neuronal survival, such as MTT assay, fractions of live/dead cells, and the extent of surviving MAP2+ neurons, revealed the protective effects of hiPSC-NSC-EVs against Aβ-42o induced neurodegeneration.

**Figure 4:**
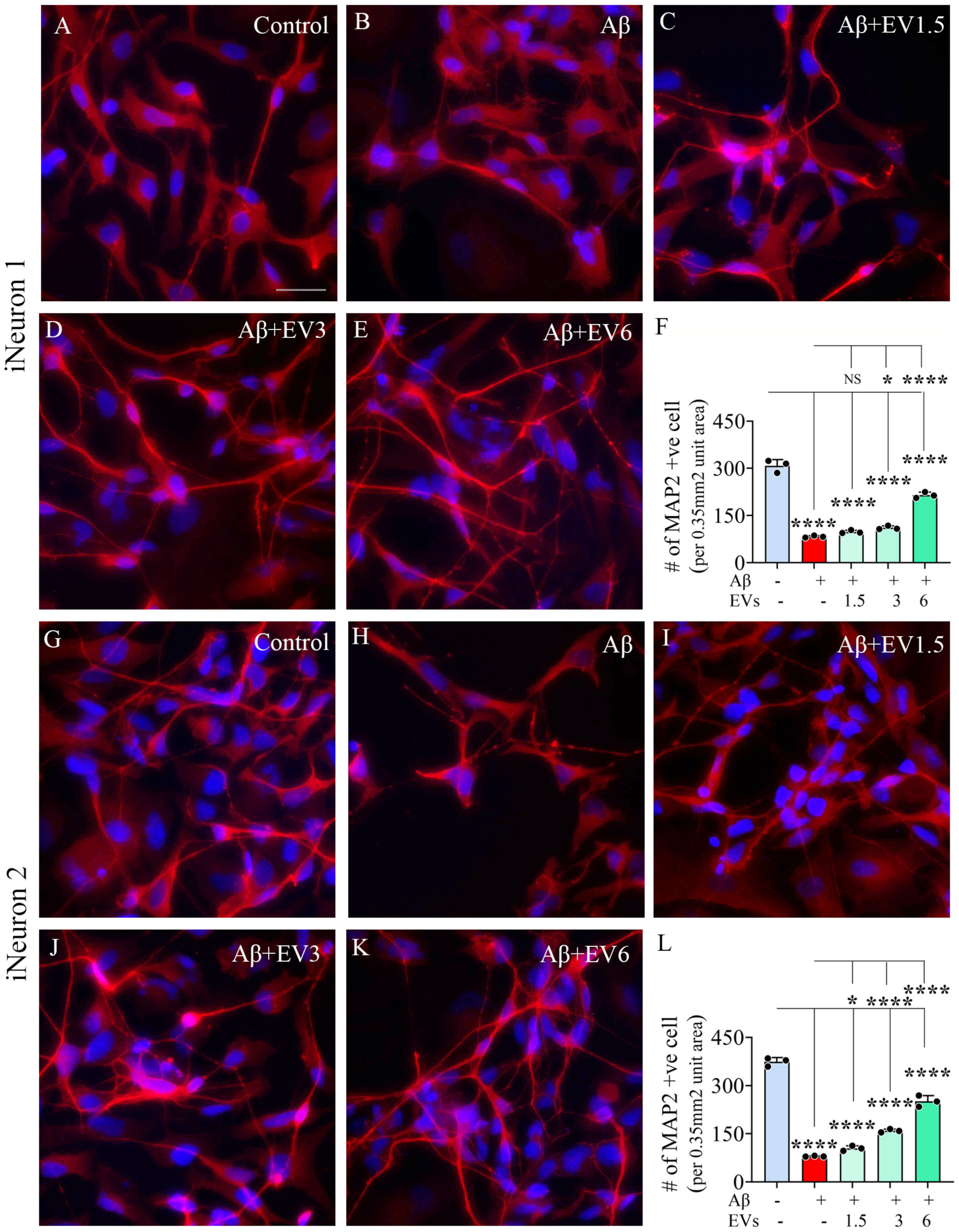
The effects of adding extracellular vesicles from human induced pluripotent stem cell-derived neural stem cells (hiPSC-NSC-EVs) into iNeurons cultures on Aβ-42 oligomers (Aβ-42o)- induced neurodegeneration. Figures A-E and H-L show representative images of surviving microtubule-associated protein-2 (MAP-2) positive neurons from control cultures of iNeurons 1 and 2 (A, G), and iNeurons 1 and 2 cultures exposed to Aβ-42o alone (B, H), Aβ-42o and treated with 1.5 x 10^9^ billion EVs (C, I), Aβ-42o and treated with 3.0 x 10^9^ billion EVs (D, J), or Aβ-42o and treated with 6.0 x 10^9^ billion EVs (E, K). Scale bar, 100 µm. The bar charts in F and L compare the numbers of surviving MAP-2+ neurons per unit area (0.35mm^2^) of iNeurons 1 and iNeurons 2 cultures across various groups.

### hiPSC-NSC-EVs treatment suppressed oxidative stress in human neurons exposed to Aβ-42o

Adding Aβ-42o to mature human neuronal cultures increased oxidative stress in neurons. In iNeurons 1 and iNeurons 2 cultures, the concentrations of oxidative stress markers MDA and PCs were elevated in cultures exposed to Aβ-42o compared to control cultures (p<0.05-0.001, Fig. 5 [A-D]). The addition of hiPSC-NSC-EVs to Aβ-42o exposed neuronal cultures dose-dependently decreased the concentrations of MDA and PCs. In iNeurons 1 cultures, compared to cultures exposed to Aβ-42o alone, MDA concentration was reduced with 1.5, 3.0, or 6 x 10^9^ doses (p<0.05-0.0001, Fig. 5 [A]), and PCs were reduced with the 6 x 10^9^ dose (p<0.05, Fig. 5 [B]). Moreover, the concentrations of PCs in EVs treated cultures were normalized to control culture levels with all doses of EVs tested (p>0.05, Fig. 5 [B]). In iNeurons 2 cultures, compared to cultures exposed to Aβ-42o alone, MDA concentration was reduced with 3.0 or 6 x 10^9^ doses (p<0.05, Fig. 5 [C]), and PCs were reduced with the 3, or 6 x 10^9^ dose (p<0.05, Fig. 5 [D]). Furthermore, the concentrations of both MDA and PCs in EVs treated cultures were normalized to control culture levels with all doses of EVs tested (p>0.05, Fig. 5 [C-D]).

**Figure 5:**
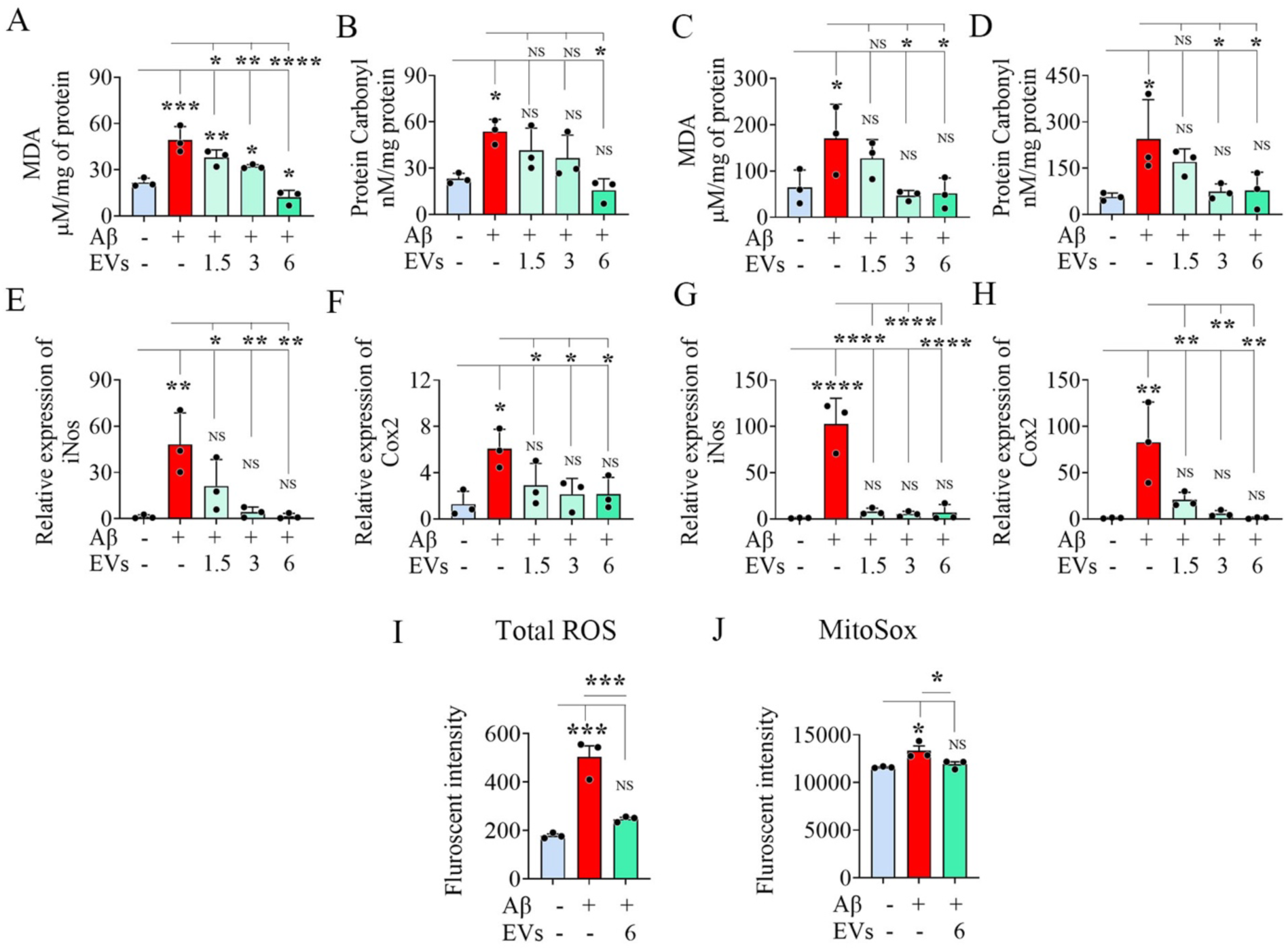
The effects of adding extracellular vesicles from human induced pluripotent stem cell-derived neural stem cells (hiPSC-NSC-EVs) into iNeurons cultures on Aβ-42 oligomers (Aβ-42o) - induced oxidative stress markers, total ROS and mitochondria-generated superoxide. The bar charts A-D compare the concentrations of malondialdehyde (MDA, A, C) and protein carbonyls (B, D) in Aβ-42o exposed cultures of iNeurons 1 (A-B) and iNeurons 2 (C-D) treated with different concentrations of hiPSC-NSC-EVs. The bar charts E-H compare the relative gene expression of iNos (E, G) and Cox2 (F, H) in Aβ-42o exposed cultures of iNeurons 1 (E-F) and iNeurons 2 (G-H) treated with different concentrations of hiPSC-NSC-EVs. The bar charts I-J compare the fluorescence intensity values for total ROS (I) and mitochondria-generated superoxide measured via MitoSOX (J) in iNeurons 2 cultures across study groups.

Next, we measured the expression of genes iNOS and COX-2 in neuronal cultures exposed to Aβ-42o. The genes iNOS and COX-2 encode proteins that are critical enzymes involved in generating ROS. The expression of these genes was increased in neuronal cultures exposed to Aβ-42o alone (p<0.05-0.0001, Fig. 5 [E-H]). In both iNeurons 1 and iNeurons 2 cultures, compared to cultures exposed to Aβ-42o alone, the expression of iNOS and COX-2 genes was reduced with all doses of EVs (1.5, 3.0 or 6 x 10^9^) tested in the study (p<0.05-0.0001, Fig. 5 [E-H]). Furthermore, their expression was normalized to control culture levels with all doses of EVs tested (p>0.05, Fig. 5 [E-H]). Thus, measurement of MDA, PCs, and genes iNOS and COX-2 all pointed to reduced oxidative stress in Aβ-42o exposed neuronal cultures treated with hiPSC-NSC-EVs.

We also measured the total ROS and MitoSOX levels in the iNeurons 1 cultures exposed to Aβ-42o. The total ROS concentration was increased in neuronal cultures exposed to Aβ-42o alone, compared to control cultures (p<0.001, Fig.5 [I]). Similarly, the concentration of mitochondria-generated superoxide measured via MitoSOX assay revealed increased levels in Aβ-42o treated cultures compared to control cultures (p<0.05, Fig.5 [J]). Notably, the levels of both total ROS and mitochondrial superoxide were normalized to levels in control cultures in Aβ-42o exposed neuronal cultures receiving hiPSC-NSC-EVs (p>0.05, Fig.5 [I-J]).

### hiPSC-NSC-EVs treatment mediated anti-apoptotic effects on human neurons exposed to Aβ-42o

The expression of pro-apoptotic genes Bad and Bax was increased in human neuronal cultures exposed to Aβ-42o. The increases were significant for both Bad and Bax in iNeurons 1 cultures and for only Bax in iNeurons 2 cultures (p<0.01, Fig. 6 [A-B, D]). In iNeurons 1 cultures, compared to cultures exposed to Aβ-42o alone, the expression of Bad and Bax genes was reduced with all doses of EVs (1.5, 3.0 or 6 x 10^9^) tested in the study (p<0.05, Fig. 6 [A-B]). In iNeurons 2 cultures, compared to cultures exposed to Aβ-42o alone, the expression of the Bad gene was reduced with 3.0 or 6 x 10^9^ EVs, whereas the expression of the Bax gene was reduced with all doses of EVs tested (p<0.05-0.01, Fig. 6 [C-D]). Furthermore, the expression of both Bad and Bax genes was normalized to control culture levels with all doses of EVs tested (p>0.05, Fig. 6 [C-D]). Consistent with these results, expression of antiapoptotic gene Bcl2 was reduced in neuronal cultures exposed to Aβ-42o (p<0.05) in both iNeurons 1 and iNeurons 2 cultures (Fig. 6 [E, G]). In iNeurons 1 cultures, compared to cultures exposed to Aβ-42o alone, the expression of Bcl-2 gene was increased in cultures treated with hiPSC-NSC-EVs with all doses of EVs (1.5, 3.0, or 6 x 10^9^ EVs) tested in the study (p<0.05-0.0001, Fig. 6 [E]). In iNeurons 2 cultures, a similar trend was observed, but increases were not significant. However, the expression of Bcl-2 was normalized to control culture levels with all doses of EVs tested in both iNeurons 1 and iNeurons 2 cultures (p>0.05, Fig. 6 [E, G]).

**Figure 6:**
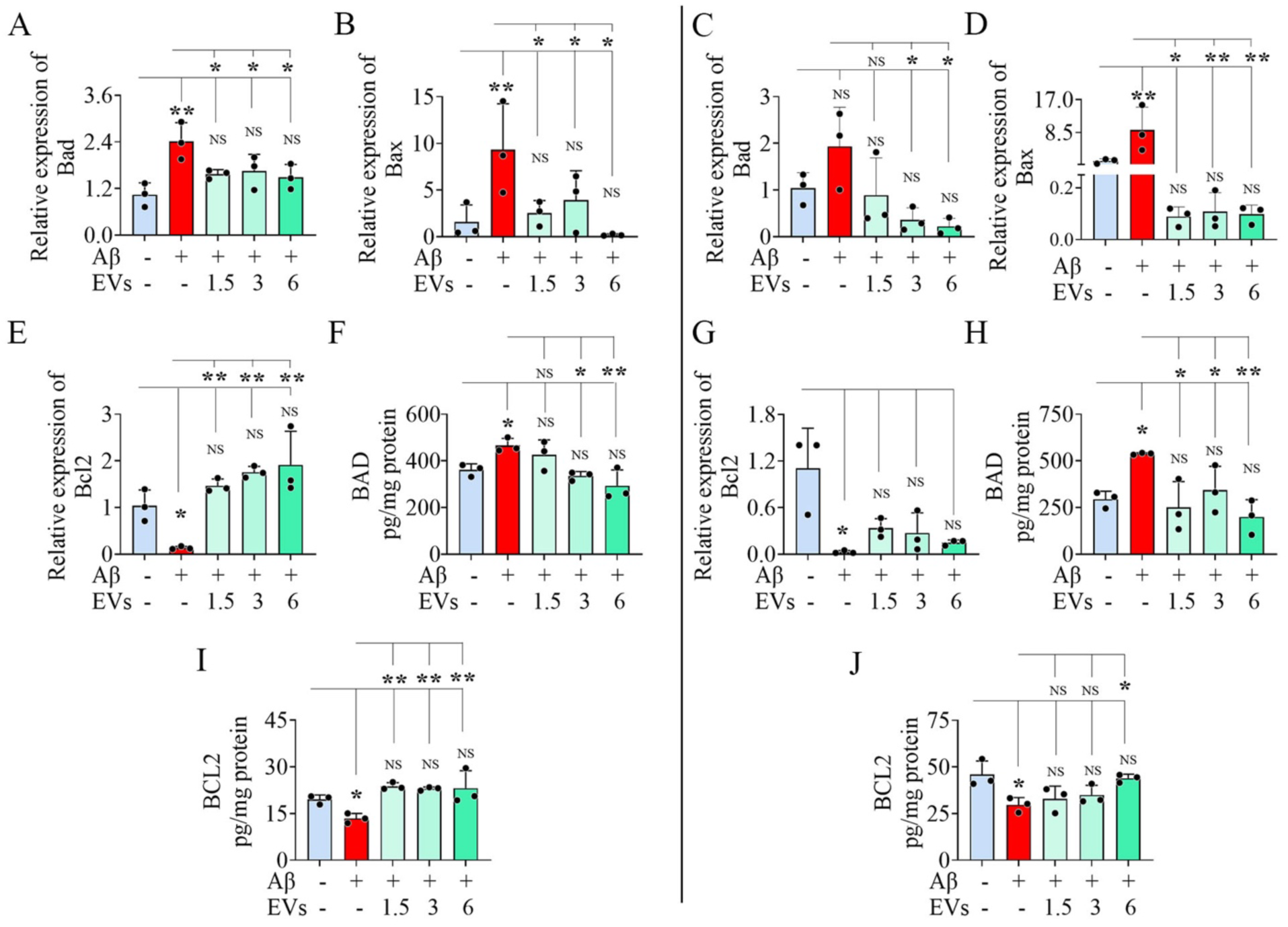
The effects of adding extracellular vesicles from human induced pluripotent stem cell-derived neural stem cells (hiPSC-NSC-EVs) into iNeurons cultures on Aβ-42 oligomers (Aβ-42o)-induced pro- and anti-apoptotic gene and protein expression. The bar charts A-E and C-G compare the relative expression of genes Bad (A, C), Bax (B, D), and Bcl2 (E, G) in Aβ-42o exposed cultures of iNeurons 1 (A-B and E) and iNeurons 2 (C-D and G) treated with different concentrations of hiPSC-NSC-EVs. The bar charts F, H, I, and J compare the protein concentrations of BAD (F, H) and BCL2 (I, J) in Aβ-42o exposed cultures of iNeurons 1 (F, I) and iNeurons 2 (H, J) treated with different concentrations of hiPSC-NSC-EVs.

We also measured the concentration of BAD and BCL2 proteins to confirm gene expression results, which revealed similar findings. In both iNeurons 1 and iNeurons 2 cultures, compared to controls, the concentration of BAD was increased in cultures exposed to Aβ-42o alone (p<0.05) but not in cultures treated with Aβ-42o and different doses of hiPSC-NSC-EVs (p>0.05, Fig. 6 [F, H]). Compared to cultures exposed to Aβ-42o alone, the concentration of BAD was decreased with 3.0 or 6 x 10^9^ EVs in iNeurons 1 cultures and with all doses of EVs tested in iNeurons 2 cultures (p<0.05-0.01, Fig. 6 [F, H]). In both iNeurons 1 and iNeurons 2 cultures, compared to controls, the concentration of BCL2 was decreased in cultures exposed to Aβ-42o alone (p<0.05) but not in cultures treated with Aβ-42o and different doses of hiPSC-NSC-EVs (p>0.05, Fig. 6 [I, J]). Compared to cultures exposed to Aβ-42o alone, the concentration of BCL2 was increased with all doses of EVs tested 3.0 or 6 x 10^9^ EVs in iNeurons 1 cultures and with 6 x 10^9^ EVs in iNeurons 2 cultures (p<0.05-0.01, Fig. 6 [I, J]). Thus, measurement of genes Bad, Bax, and Bcl-2 and proteins BAD and BCL2 supported significant antiapoptotic effects of hiPSC-NSC-EVs on neuronal cultures exposed to Aβ-42o.

### hiPSC-NSC-EVs treatment improved the mitochondrial membrane potential in human neurons exposed to Aβ-42o

JC-1 assay followed by fluorimetry reading revealed a high ratio of red fluorescence in control human neuronal cultures, implying a high mitochondrial membrane potential and healthy mitochondria (Fig. 7 [A, D, G, J]). Adding Aβ-42o to neuronal cultures resulted in a lower ratio of red fluorescence, with a concomitant increase in green fluorescence intensity, suggesting a lower mitochondrial potential or damaged mitochondria (Fig. 7 [B, E, H, J], p<0.01 compared to control cultures). However, in Aβ-42o exposed neuronal cultures receiving hiPSC-NSC-EVs, the ratio of red fluorescence improved (Fig. C, F, I, J], p<0.05 compared to cultures exposed to Aβ-42o alone). AF analysis of red/green ratio from images using ImageJ revealed that Aβ-42o exposed neuronal cultures exhibited a reduced ratio of red fluorescence than control neuronal cultures (p<0.05), and Aβ-42o exposed neuronal cultures receiving hiPSC-NSC-EVs displayed a similar ratio of red fluorescence as control neuronal cultures (Fig. 7 [K]).

**Figure 7:**
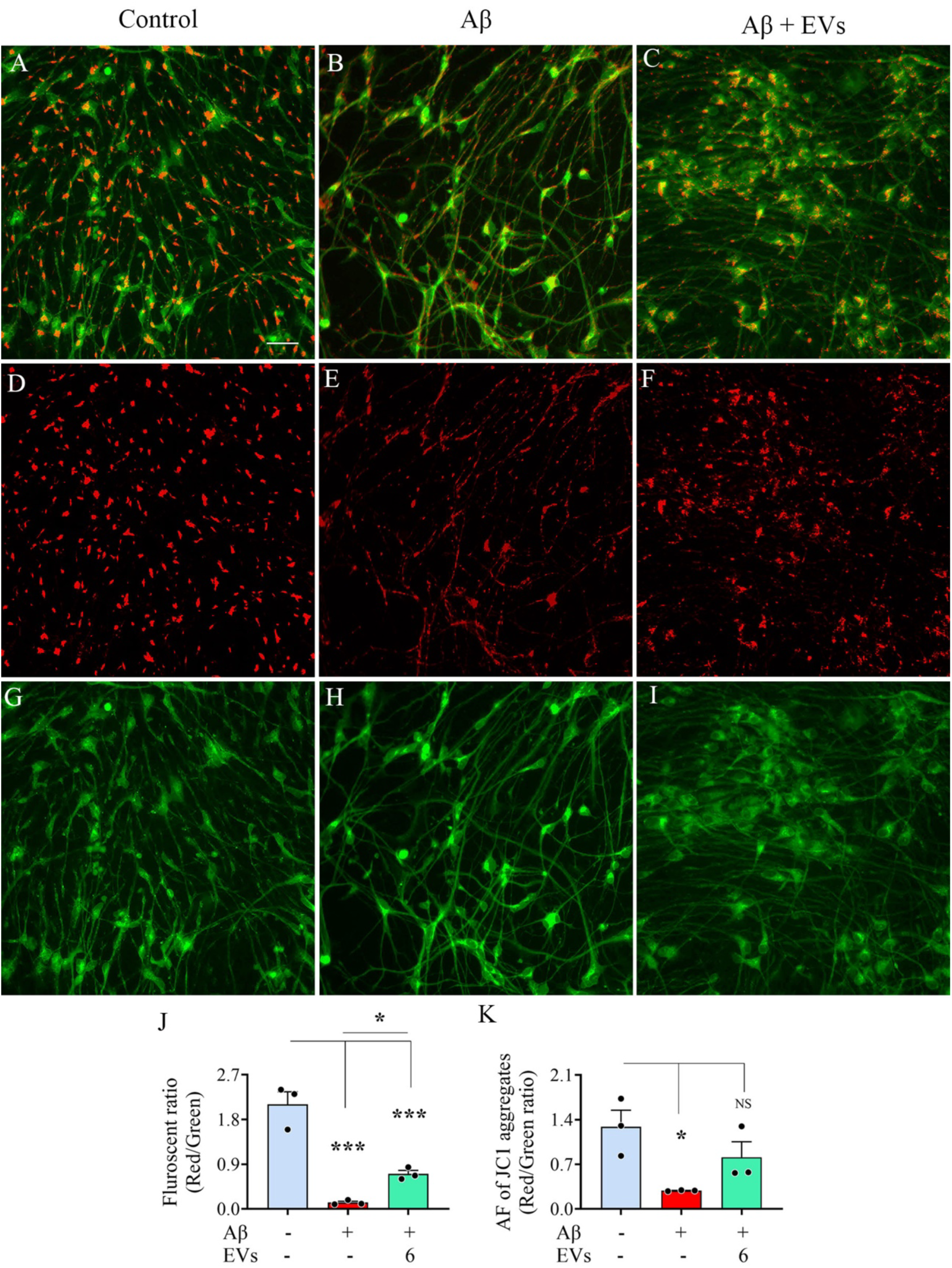
The effects of adding extracellular vesicles from human induced pluripotent stem cell-derived neural stem cells (hiPSC-NSC-EVs) into iNeurons cultures on Aβ-42 oligomers (Aβ-42o)-induced impairment of mitochondrial membrane potential. Figures A-I are representative images of red and green fluorescence from a JC-1 assay in neuronal cultures from a control group (A, D, G), Aβ-42o treated group (B, E, H), or Aβ-42o and 6 x 10^9^ hiPSC-NSC-EVs treated group (C, F, I). The bar charts compare the ratio of red/green fluorescence measured through fluorimetry (J) or area fraction (AF) analysis using Image J (K) across control, Aβ-42o exposed, and Aβ-42o and 6 x 10^9^ hiPSC-NSC-EVs treated groups. Scale bar, 100 µm.

### Effect of hiPSC-NSC-EVs treatment on total mitochondria in human neurons exposed to Aβ-42o

The mitochondrial outer membrane staining with the TOM20 antibody suggested reduced mitochondria in the soma and dendrites of neurons of cultures exposed to Aβ-42o alone compared to control and hiPSC-NSC-EVs-treated cultures (Fig. 8 [A-I] and Supplementary Fig. 6). Comparison of the average AF of TOM20+ structures within the soma of individual neurons between groups revealed a significantly decreased amount of mitochondria in human neuronal cultures treated with Aβ-42o alone, compared to control cultures (p<0.01, Fig. 8 [J]). However, in human neuronal cultures exposed to Aβ-42o receiving hiPSC-NSC-EVs, the average amount of TOM20+ mitochondria per neuron was maintained at the control culture level (p>0.05, Fig. 8 [J]). It was also higher than in cultures exposed to Aβ-42o alone (p<0.05, Fig. 8 [J]). Thus, hiPSC-NSC-EVs can prevent mitochondria loss in human neurons in response to Aβ-42o exposure.

**Figure 8:**
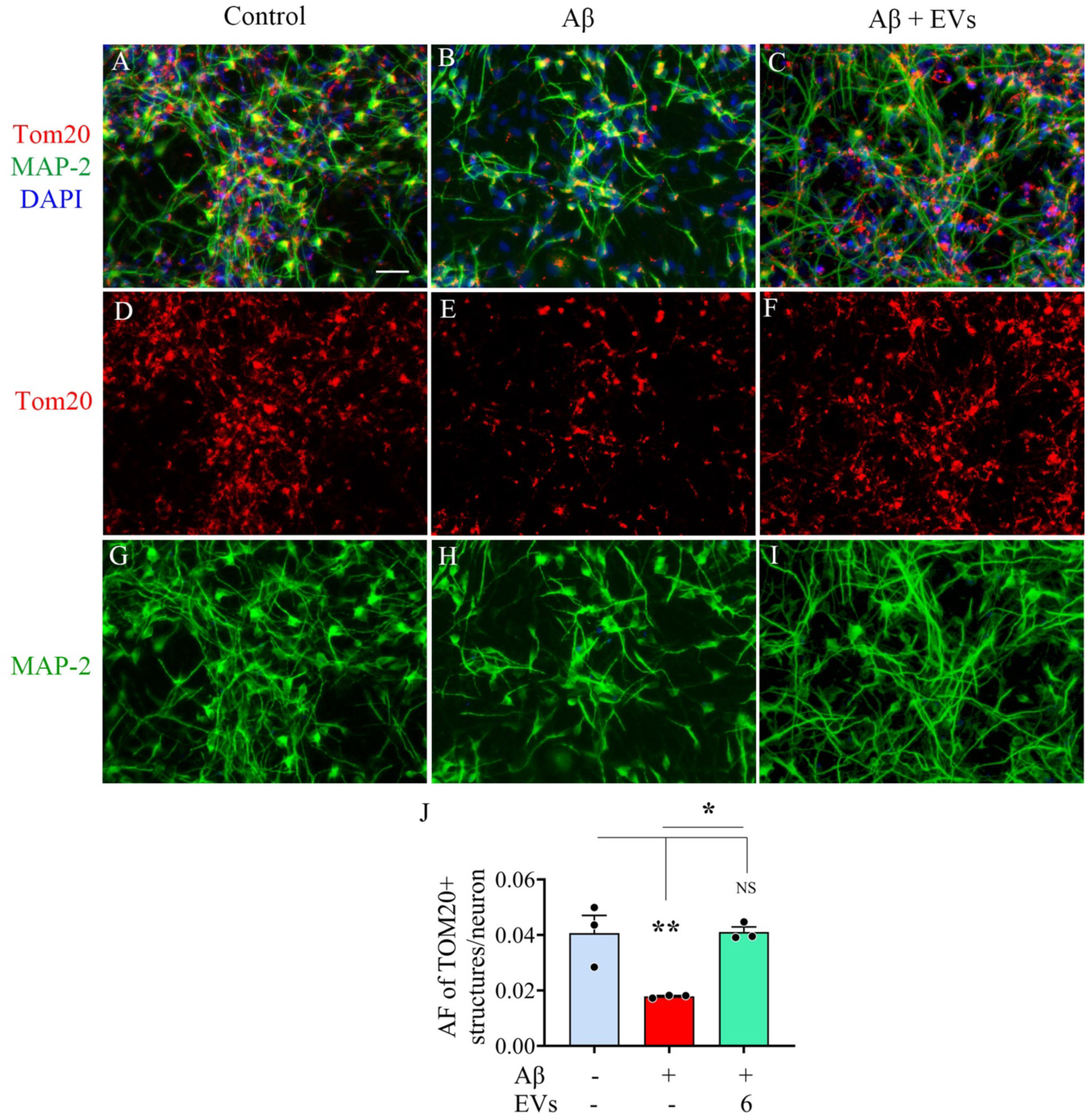
The effects of adding extracellular vesicles from human induced pluripotent stem cell-derived neural stem cells (hiPSC-NSC-EVs) into iNeurons cultures on Aβ-42 oligomers (Aβ-42o)-induced mitochondrial depletion. Figures A-I show representative images of TOM20+ structures (red, visualizing mitochondria) in Map-2+ neurons (green) in cultures from a control group (A, D, G), Aβ-42o treated group (B, E, H), or Aβ-42o and 6 x 10^9^ hiPSC-NSC-EVs treated group (C, F, I). The bar charts compare the area fraction (AF) TOM20+ mitochondria measured by Image J (J) across control, Aβ-42o exposed, and Aβ-42o and 6 x 10^9^ hiPSC-NSC-EVs treated groups. Scale bar, 100 µm.

### hiPSC-NSC-EVs treatment normalized the expression of genes linked to mitochondrial complex proteins in human neurons exposed to Aβ-42o

The expression of genes linked to mitochondrial complex such as Ndufs6, Ndufs7 (complex I), Sdha, Sdhb (Complex II), and Atp6ap1 (complex V) were upregulated in neuronal cultures exposed to Aβ-42o (p<0.05-0.01, Fig. 9 [A-D, F, H]). However, in Aβ-42o exposed neuronal cultures treated with 6 x 10^9^ EVs, their expression levels were normalized to naïve control levels (p>0.05, Fig 9 [A-D, F, H]). Furthermore, compared to neuronal cultures exposed to Aβ-42o alone, Aβ-42o exposed neuronal cultures treated with 6 x 10^9^ EVs displayed reduced expression of Ndufs6, Ndufs7, Sdha, Sdhb, and Atp6ap1 (p<0.05-0.01, Fig 9 [A-D, F, H]). The expression of genes Cyc1, Bcs1l (Complex III), and Cox6a1 (complex IV) also showed similar trends but were not significant statistically. Thus, genes linked to the mitochondrial electron transport undergo downregulation in human neurons exposed to Aβ-42o, but their expression could be normalized through hiPSC-NSC-EVs treatment.

**Figure 9:**
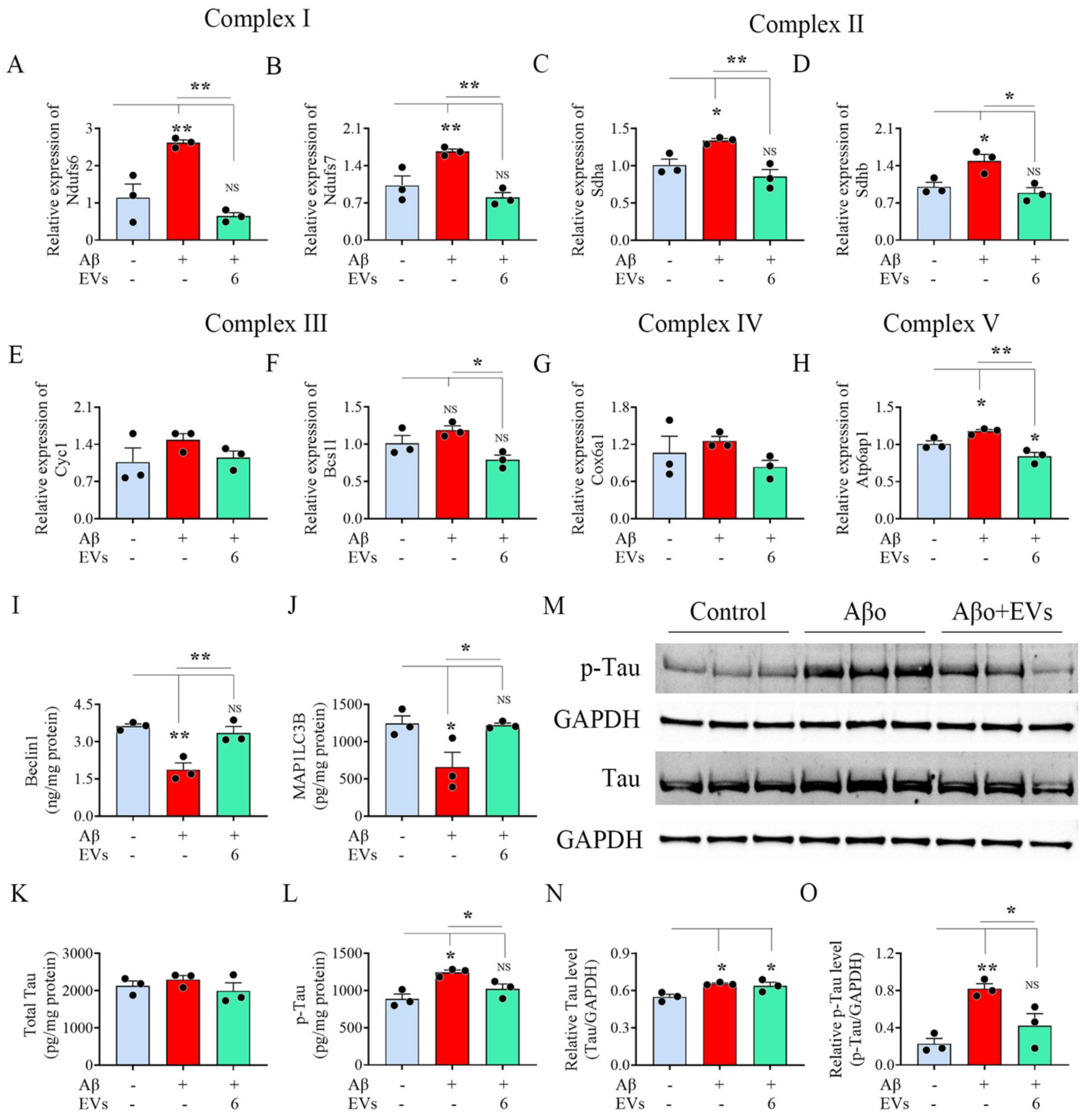
The effects of adding extracellular vesicles from human induced pluripotent stem cell-derived neural stem cells (hiPSC-NSC-EVs) into iNeurons cultures on Aβ-42 oligomers (Aβ-42o)-induced increase in the expression of genes encoding mitochondrial complex proteins, decrease in autophagy-related proteins and tau phosphorylation. The bar charts A-H compare the relative expression of genes Ndufs6 (A), Ndufs7 (B), Sdha (C), Sdhb (D), Cyc1 (E), Bcs1l (F), Cox6a1 (G) and Atp6ap1 (H) in neuronal cultures (iNeurons 1) between control, Aβ-42o exposed, and Aβ-42o and 6 x 10^9^ hiPSC-NSC-EVs treated groups. The bar charts I and J compare the concentrations of autophagy-related proteins Beclin 1 (I) and MAP1LC3B (J) in neuronal cultures (iNeurons 1) between control, Aβ-42o exposed, and Aβ-42o and 6 x 10^9^ hiPSC-NSC-EVs treated groups. The bar charts K and L compare the concentrations of total tau (K) and phosphorylated-tau (p-tau, L) neuronal cultures (iNeurons 1) between control, Aβ-42o exposed, and Aβ-42o and 6 x 109 hiPSC-NSC-EVs treated groups. The blot M displays the relative tau and p-tau levels in neuronal cultures (iNeurons 1) from the control, Aβ-42o-treated, and Aβ-42o plus 6 x 10^9^ hiPSC-NSC-EVs treated groups. The bar charts N and O compare the relative levels of total tau (K) and phosphorylated tau (p-tau, L) in neuronal cultures between control, Aβ-42o exposed, and Aβ-42o and 6 x 10^9^ hiPSC-NSC-EVs treated groups.

### hiPSC-NSC EVs treatment improved autophagy in human neurons exposed to Aβ-42o

The concentration of autophagy-related proteins Beclin-1 and MAP1-LC3B were downregulated in human neuronal cultures exposed to Aβ-42o (p<0.05-0.01, Fig. 9 [I-J]). However, in Aβ-42o exposed neuronal cultures treated with 6 x 10^9^ EVs, their concentrations were normalized to naïve control levels (p>0.05, Fig 9 [I-J]). Furthermore, compared to neuronal cultures exposed to Aβ-42o alone, Aβ-42o exposed neuronal cultures treated with 6 x 10^9^ EVs displayed increased concentration of Beclin-1 and MAP1-LC3B (p<0.05-0.01, Fig 9 [I-J]). Thus, hiPSC-NSC-EVs treatment can normalize autophagy in human neurons exposed to Aβ-42o.

### hiPSC-NSC EVs treatment reduced increased tau phosphorylation in human neurons exposed to Aβ-42o

The concentration of total tau did not change in human neurons exposed to Aβ (Fig. 9 [K]), but the phosphorylation of tau increased with Aβ-42o exposure (Fig. 9 [L]). The concentration of p-tau increased in neuronal cultures exposed to Aβ-42o (p<0.05, Fig. 9 [L]). However, in Aβ-42o exposed neuronal cultures treated with 6 x 10^9^ EVs, the concentration of p-tau was normalized to the naïve control level (p>0.05, Fig 9 [L]). Furthermore, compared to neuronal cultures exposed to Aβ-42o alone, Aβ-42o exposed neuronal cultures treated with 6 x 10^9^ EVs displayed reduced concentration of p-tau (p<0.05, Fig. 9 [L]). Western blot analysis further supported these results (Fig.9 [M]). The total tau and p-tau levels were significantly increased in human neuronal cultures exposed to Aβ-42o alone compared to control cultures (p<0.05-0.01, Fig. 9 [N-O]). However, in human neuronal cultures exposed to Aβ-42o receiving hiPSC-NSC-EVs, while the total tau level was higher than in control cultures, the extent of p-tau was normalized to the naive control level (p>0.05, Fig. 9 [O]). It was also lower than in cultures exposed to Aβ-42o alone (p<0.05, Fig. 9 [O]). Thus, human neurons exposed to Aβ-42o undergo increased tau phosphorylation but hiPSC-NSC-EVs can considerably reduce such adverse change.

## Discussion

The findings of this study underscore the potential of purified and well-characterized hiPSC-NSC-EVs in safeguarding human neurons derived from two distinct hiPSC lines against Aβ-42o induced neurodegeneration. In the context of Aβ-42o induced neurodegeneration, a moderate dose of 1 µM Aβ-42o was selected for neuroprotection experiments, as higher doses were observed to induce more severe neurodegeneration in a dose-dependent manner. The prominent pathological alterations induced by 1 µM Aβ-42o in human neurons encompassed increased production of total ROS and mitochondria-generated superoxide, diminished mitochondrial membrane potential, heightened lipid peroxidation, and protein oxidation. Additionally, there was an augmented expression of iNOS and COX-2 genes that encode proteins implicated in ROS generation, increased expression of pro-apoptotic genes and proteins (Bad and Bax), alongside decreased expression of an antiapoptotic gene and protein (Bcl-2). Furthermore, loss and hyperactivation of the mitochondria were evident from reduced TOM20+ structures and the upregulation of genes associated with mitochondrial complexes, including Ndufs6, Ndufs7 (complex I), Sdha, Sdhb (Complex II), and Atp6ap1 (complex V). Moreover, exposure of human neurons to 1 µM Aβ-42o appeared to influence autophagy, as it reduced concentrations of autophagy-related proteins Beclin-1 and MAP1-LC3B. Additionally, Aβ-42o-exposed human neurons displayed increased tau phosphorylation. Remarkably, introducing hiPSC-NSC-EVs into Aβ-42o-exposed human neuronal cultures provided dose-dependent protection to human neurons through antioxidant and antiapoptotic mechanisms. These protective effects were evidenced by reduced levels of total ROS, mitochondria-derived superoxide, higher mitochondrial membrane potential, diminished concentrations of MDA and PCs, reduced expression of iNOS, COX-2, Bad, and Bax, increased expression of Bcl-2 and TOM20+ mitochondria, and the normalized expression of genes involved in mitochondrial complex proteins and autophagy. These effects were also associated with the mitigation of tau phosphorylation.

### hiPSC-NSC-EVs protected human neurons exposed to Aβ-42o by antioxidant activity and normalizing mitochondrial function

Previous research has reported that Aβ-42o can induce pathological changes and toxicity in rodent neurons. Studies have indicated that Aβ42o can negatively impact Ca^2+^ homeostasis in neurons by promoting increased Ca^2+^ entry [61]. Depending on the dosage, this effect may result in a significant Ca^2+^ influx and excitotoxicity [62,63]. Aβ-42o is also able to integrate into the lipid bilayer, acting as a source of ROS and initiating lipid peroxidation [23–25]. This membrane lipid peroxidation can disrupt neuronal Ca^2+^ homeostasis, leading to Ca^2+^ overload and neurodegeneration [64]. Studies have also shown that Aβ-42o can form pore-like structures (also referred to as amyloid channels) in neuronal membranes causing increased Ca^2+^ permeability, disruption of Ca^2+^ homeostasis, and eventually neurodegeneration [61,65]. Furthermore, Aβ-42o could accumulate in mitochondrial membranes, leading to a reduction in mitochondrial membrane potential. JC-1 assay in this study confirmed reduced mitochondrial membrane potential in human neurons with Aβ-42o exposure. Accumulation Aβ-42o also disrupts the transportation of nuclear-encoded mitochondrial proteins to mitochondria, interferes with the electron transport chain, increases the production of ROS, and results in damage to the mitochondria [26]. Indeed, total ROS and MitoSOX assays in this study validated increased ROS and mitochondria-derived superoxide production in human neurons with Aβ-42o exposure. Moreover, exposure of neurons to Aβ-42o can lead to a decrease in mitochondrial superoxide dismutase 2 and mitochondrial cytochrome C protein levels [27]. While the current study did not explore altered Ca^2+^ homeostasis, it did observe increased lipid peroxidation in human neurons exposed to Aβ-42o, as evidenced by elevated MDA concentration. An overall increase in oxidative stress was also apparent from enhanced levels of PCs and iNOS. iNOS is expressed only during pathophysiological states, and the nitric oxide produced by iNOS enhances the production of ROS [66]. The elevated expression of Cox-2 also indicated increased oxidative stress in human neurons exposed to Aβ-42o. Neurons are particularly susceptible to damage caused by free radicals generated through COX-2 peroxidase activity [67].

Notably, the addition of hiPSC-NSC-EVs into Aβ-42o-exposed human neuronal cultures protected them by significantly reducing total ROS and superoxide production, lipid peroxidation, as evidenced by decreased MDA levels, and overall oxidative stress, as observed from the reduced expression of PCs, iNOS, and COX-2. These antioxidant effects of hiPSC-NSC-EVs may be attributed to specific miRNAs and protein cargo they carry. For example, hiPSC-NSC-EVs are enriched with hemopexin, which can maintain mitochondrial function and homeostasis, as well as reducing ROS toxicity [36,44]. Indeed, mitochondrial dysfunction in human neurons induced by Aβ-42o exposure was prevented by hiPSC-NSC-EVs treatment in this study. Such modulation in human neurons exposed to Aβ-42o and hiPSC-NSC-EVs could be gleaned from a higher mitochondrial membrane potential, normalized expression of genes linked to mitochondrial complexes I (Ndufs6, Ndufs7), II (Sdha, Sdhb), and IV (Atp6ap1) to levels seen in naive human neurons and maintenance of higher levels of mitochondria. Thus, antioxidant and mitochondria protecting effects are among the mechanisms by which hiPSC-NSC-EVs protected human neurons exposed to Aβ-42o.

### hiPSC-NSC-EVs protected human neurons exposed to Aβ-42o by antiapoptotic effects

Exposure of neurons to Aβ-42o results in apoptosis, which involves altered expression of pro- and anti-apoptotic proteins [29]. Particularly, mitochondrial-specific accumulation of Aβ-42o induces mitochondrial dysfunction causing apoptotic cell death [28]. Indeed, in this study, the neurodegeneration occurring through apoptosis was evident from increased expression of pro-apoptotic genes and proteins Bad and Bax, alongside decreased expression of an antiapoptotic gene and protein (Bcl-2). This is consistent with the Bax and Bad upregulation and Bcl-2 downregulation found in hippocampal slice cultures exposed to Aβ-42o [68] and altered expression of these proteins within neurons in AD [69]. Bax promotes apoptosis by translocating into the mitochondrial membrane and facilitating cytochrome c release whereas Bad induces the activation of Bax, by inactivating the pro-survival Bcl-2 protein. Bcl-2 prevents apoptotic death in multiple cell types including neurons [70]. Bcl-2 prevents the permeabilization of mitochondria to inhibit Ca^2+^ overload and consequently the activity of cleaved caspases, whereas Bax exerts the opposite effects [71].

However, the extent of apoptosis was reduced after introducing hiPSC-NSC-EVs into Aβ-42o-exposed human neuronal cultures. Such a reduction could be ascertained from the decreased expression of Bad and Bax and increased expression of Bcl-2. In this study, the downregulation of Bax by hiPSC-NSC-EVs is likely the most crucial antiapoptotic mechanism, as previous study has shown that the suppression of Aβ-42o-mediated neurotoxicity can be significantly achieved through the inhibition of Bax activity, either genetically or pharmacologically [68]. Additionally, the antiapoptotic effects of hiPSC-NSC-EVs are likely due to miRNA-103a-3p and miRNA-21-5p, as well as PTX3 protein carried by them, as they can promote neuroprotection through antiapoptotic mechanisms. [39–42]. Our study has validated the presence of these miRNAs and PTX3 in hiPSC-NSC-EVs employed in the study. The role of miRNA-103a in inhibiting apoptosis has been observed in several studies. One study demonstrated that during hypoxia/reoxygenation conditions, overexpression of miRNA-103a in cardiomyocytes significantly reduced apoptosis by inhibiting Bax, cleaved caspase-3, and cleaved caspase-9 [72]. Another study showed that in nerve growth factor-stimulated PC12 cells and primary cerebral cortical neurons exposed to Aβ-42o, overexpression of miRNA-103 resulted in decreased apoptosis associated with reduced expression of cleaved caspase-3, while inhibition of miRNA-103 led to increased apoptosis [42]. Further analysis indicated that miRNA-103 exerts antiapoptotic effects by targeting prostaglandin-endoperoxide synthase 2, a protein that plays a significant role in AD development and progression [42]. Thus, hiPSC-NSC EVs enriched with miRNA-103a have the potential for mitigating neurodegeneration in AD. This is particularly important given that miRNA-103 is downregulated in AD patients and models [73,74], indicating its involvement in the pathogenesis of AD.

Additionally, miRNA-21-5p, found in abundance in hiPSC-NSC-EVs, likely also played a role in mitigating apoptosis because an earlier investigation using SH-SY5Y cells showed that miRNA-21 can significantly diminish apoptosis induced by Aβ-42o. Such effect was associated with the inhibition of Bax and programmed cell death protein 4, alongside an elevation in Bcl-2 [54]. Furthermore, the contribution of pentraxin-3 protein within hiPSC-NSC-EVs to antiapoptotic properties cannot be ruled out since a previous study utilizing a mouse model of Parkinson’s disease revealed that treatment with human recombinant pentraxin-3 can prevent apoptosis and degeneration of dopaminergic neurons [55]. Considering these findings, it is conceivable that the enrichment of miRNA-103a, miRNA-21-5p, and pentraxin-3 in hiPSC-NSC-EVs has significantly attenuated apoptosis in human neurons exposed to Aβ-42o in this study. However, establishing a definitive cause-and-effect relationship would necessitate further investigations utilizing naïve EVs loaded with these miRNAs or pentraxin-3.

### hiPSC-NSC-EVs reduced tau phosphorylation and improved autophagy in Aβ-42o exposed human neurons

Aβ-42o exposures to neurons elicit a substantial increase in tau protein phosphorylation. For instance, the administration of Aβ-42o into the brains of mice induces a marked rise in the number of NFTs proximal to the injection sites [75]. Therefore, it is likely that Aβ-42o operates upstream of tau, instigating a cascade that culminates in tau-dependent synaptic dysfunction [76,77]. Aβ-42o exposure augments tau phosphorylation through the activation of tau kinases mitogen-activated protein kinase and glycogen synthase kinase-3 beta [30,31]. Furthermore, a study suggested that heightened ROS production after Aβ-42o exposure initiates mechanistic target of rapamycin C1 (mTORC1) activation [78]. Subsequent mTORC1 elevation inhibits autophagy and stimulates the expression of cell cycle regulatory proteins such as cyclin-dependent kinase 2 (CDK2), which interacts with tau, inducing tau phosphorylation and microtubule destabilization during neurodegeneration [78]. These findings align with the decreased concentration of autophagy-related proteins Beclin 1 and MAP1LC3B and increased tau phosphorylation observed in the present study.

Introducing hiPSC-NSC-EVs into Aβ-42o-exposed human neuronal cultures improved autophagy and reduced tau phosphorylation. It remains to be investigated how hiPSC-NSC-EVs improve autophagy and decrease tau phosphorylation in human neurons exposed to Aβ-42o. However, it is worth noting that hiPSC-NSC-EVs carry miRNA-148a-3p [36], which can inhibit tau phosphorylation. The potential role of miRNA-148-3a in inhibiting tau phosphorylation is evident from a study demonstrating that the downregulation of miRNA-148a-3p leads to tau hyperphosphorylation and the progression of AD [43]. Additional analysis revealed that miRNA-148a-3p targets CDK-5 regulatory subunit 1 and PTEN involved in tau hyperphosphorylation. Moreover, a strategy enhancing miRNA-148a-3p signaling reduced tau hyperphosphorylation in AD mice [43].

## Conclusions

The results of this study emphasize the potential of hiPSC-NSC-EVs in protecting human neurons derived from two distinct hiPSC lines against neurodegeneration induced by Aβ-42o. Importantly, when hiPSC-NSC-EVs were introduced into Aβ-42o-exposed human neuronal cultures, they protected human neurons in a manner that depended on the dosage. Such protection was mediated mainly through antioxidant, antiapoptotic, and mitochondria-protecting mechanisms. The evidence for such mechanisms included reduced levels of oxidative stress, proapoptotic and impaired mitochondrial function markers, and increased expression of genes and/or proteins that inhibit apoptosis. Furthermore, these beneficial effects were also associated with normalized autophagy and decreased tau phosphorylation. However, investigation of the electrophysiological properties of neurons exposed to Aβ-42o alone vis-à-vis neurons exposed to Aβ-42o and hiPSC-NSC-EVs treatment will be needed in future studies to determine whether hiPSC-NSC-EVs can prevent or alleviate the possible electrophysiological abnormalities in neurons exposed to Aβ-42o. Additionally, investigating changes in global gene expression in neurons using bulk RNA sequencing will help determine the proficiency of hiPSC-NSC-EVs in alleviating the activation of potential other pathways involved in Aβ-42o-induced neurodegeneration. Overall, the findings from this study will support investigating whether administering hiPSC-NSC-EVs in AD models would provide significant neuroprotection and slow AD pathogenesis.

## Acknowledgments

The TEM images of hiPSC-NSC-EVs were taken at the Image Analysis Laboratory, Texas A&M Veterinary Medicine & Biomedical Sciences (RRIS: SCR_0222479).

## Availability of data and materials

Data supporting this study are included within the article.

## Authors’ contributions

Concept: AKS. Research design: AKS, SR, and LNM. Data collection, analysis, and interpretation: SR, LNM, RSB, GS, SK, AN, RU, EN JJC, and AKS. Preparation of figure composites: SR, LNM and AKS. Manuscript writing: SR, LNM, and AKS. All authors provided feedback, edits, and additions to the manuscript text and approved the final version of the manuscript.

## Declarations

### Ethics approval and consent to participate

The sources of hiPSCs (LiPSC-GR1.1 and iPS (IMR90)-4) employed in this study have confirmed that ethical approval has been obtained for collecting human cells and that the donors had signed informed consent.

Additional details:

1. LiPSC-GR1.1 (Research grade), https://commonfund.nih.gov/stemcells/faq The hospital has granted permission to access medically suitable patients, and the hospital IRB has approved the informed consent document and processes necessary for donor screening and testing and tissue recovery at their site.
2. iPS (IMR90)-4, https://hpscreg.eu/cell-line/WISCi004-B Informed consent has been obtained from the donor of the embryo/tissue from which the pluripotent stem cells were derived.

### Other declarations

The authors declare that they have not used Artificial Intelligence in this study.

### Consent for publication

All necessary consent and approvals were obtained from all authors and co-authors.

