## Supplementary Figures for "Extracellular Vesicles from hiPSC-derived NSCs Protect Human Neurons against Aβ-42 Oligomers Induced Neurodegeneration, Mitochondrial Dysfunction and Tau Phosphorylation"

(1) **Supplementary Figure 1:** Western blot showing CD63 expression in hiPSC-NSC-EVs preparations.

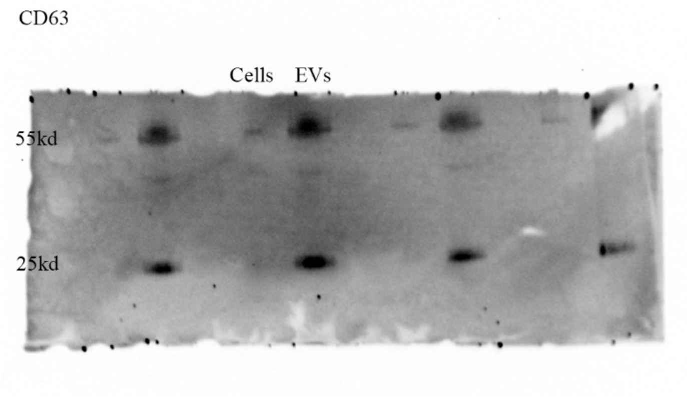


(2) **Supplementary Figure 2:** Western blot showing CD81 expression in hiPSC-NSC-EVs preparations.

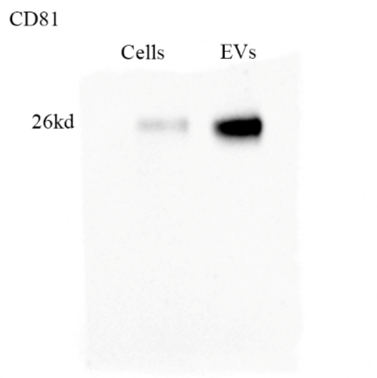


(3) **Supplementary Figure 3:** Western blot showing ALIX expression in hiPSC-NSC-EVs preparations.

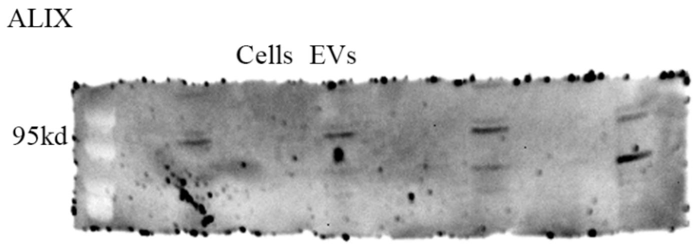

**(4) Supplementary Figure 4:** Western blot showing the absence of cytochrome C expression in hiPSC-NSC-EVs preparations.

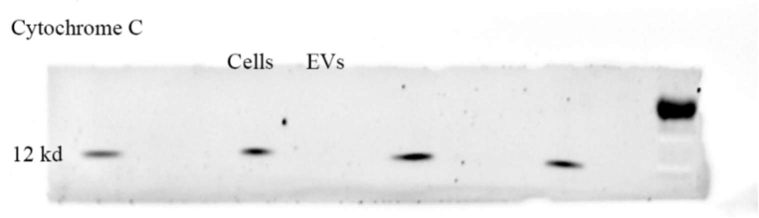


**(5) Supplementary Figure 5:** Western blots showing the relative protein levels of tau, p-tau and GAPDH in control, Aβ-42o and Aβ-42o-EVs groups.

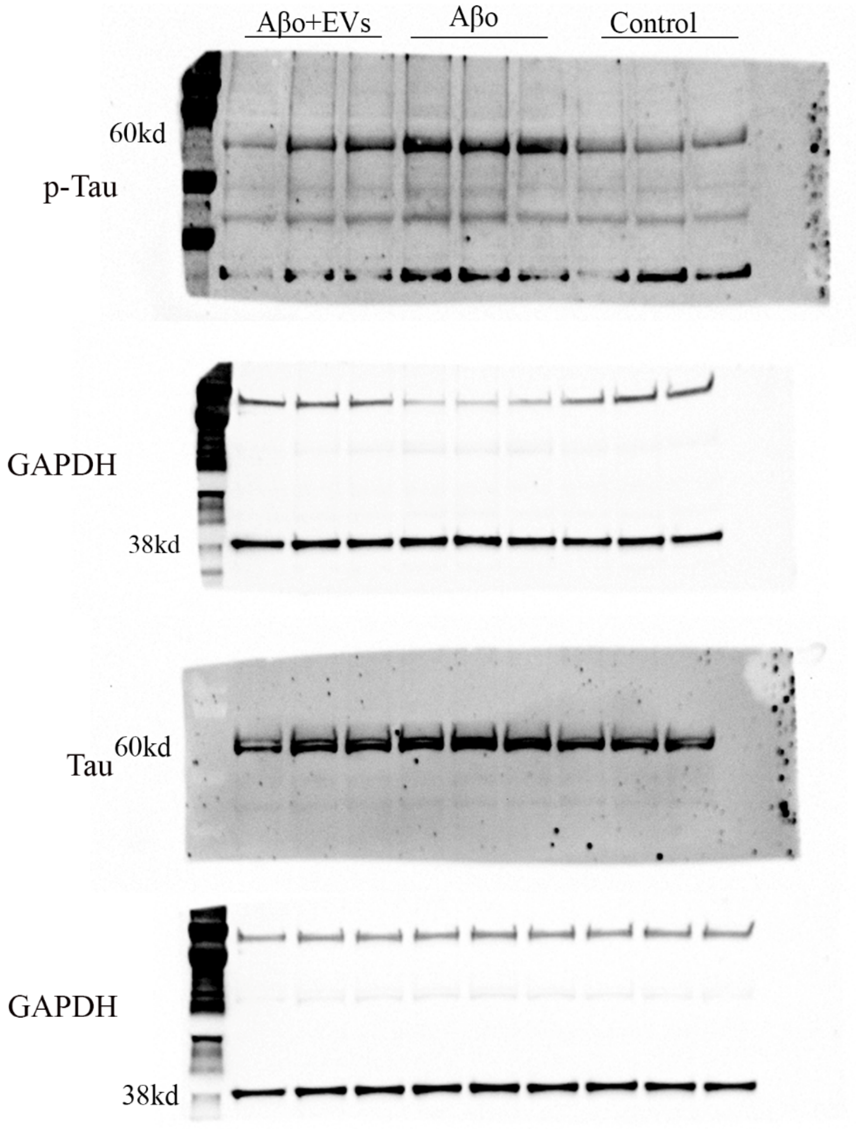

**Supplementary Figure 6**

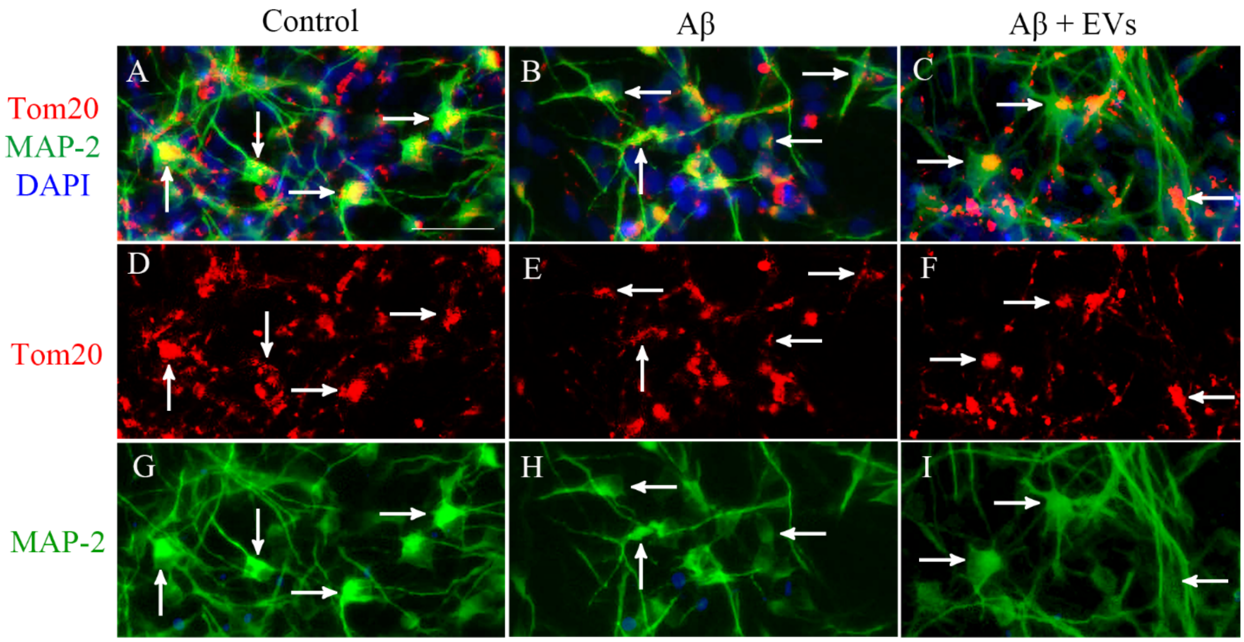


**Supplementary Figure 6:** Images showing magnified views of TOM20+ structures (red; representing mitochondria) in MAP2+ human neurons (green) from a control group (A, D, G), Aβ-42o exposed group (B, E, H), and Aβ-42o and 6 x $10^{9}$ EVs treated group (C, F, I). Scale bar, 100 µm.
